# Decreasing the flexibility of the TELSAM-target protein linker and omitting the cleavable fusion tag improves crystal order and diffraction limits

**DOI:** 10.1101/2023.05.12.540586

**Authors:** Parag L. Gajjar, Maria J. Pedroza Romo, Celeste M. Litchfield, Miles Callahan, Nathan Redd, Supeshala Nawarathnage, Sara Soleimani, Jacob Averett, Elijah Wilson, Andrew Lewis, Cameron Stewart, Yi-Jie J. Tseng, Tzanko Doukov, Andrey Lebedev, James D. Moody

**Author notes:** IMPORTANT: this document contains embedded data - to preserve data integrity, please ensure where possible that the IUCr Word tools (available from http://journals.iucr.org/services/docxtemplate/) are installed when editing this document.

## Abstract

TELSAM crystallization promises to become a revolutionary tool for the facile crystallization of proteins. TELSAM can increase the rate of crystallization and form crystals at low protein concentrations without direct contact between TELSAM polymers and, in some cases, with very minimal crystal contacts overall (Nawarathnage *et al*., 2022). To further understand and characterize TELSAM-mediated crystallization, we sought to understand the requirements for the composition of the linker between TELSAM and the fused target protein. We evaluated four different linkers Ala-Ala, Ala-Val, Thr-Val, and Thr-Thr, between 1TEL and the human CMG2 vWa domain. We compared the number of successful crystallization conditions, the number of crystals, the average and best diffraction resolution, and the refinement parameters for the above constructs. We also tested the effect of the fusion protein SUMO on crystallization. We discovered that rigidification of the linker improved diffraction resolution, likely by decreasing the number of possible orientations of the vWa domains in the crystal, and that omitting the SUMO domain from the construct also improved the diffraction resolution.

**Synopsis:** We demonstrate that the TELSAM protein crystallization chaperone can enable facile protein crystallization and high-resolution structure determination. We provide evidence to support the use of short but flexible linkers between TELSAM and the protein of interest and to support the avoidance of cleavable purification tags in TELSAM-fusion constructs.

## 1. Introduction

Experimental structural characterization of proteins is essential for protein engineering, drug and target discovery, virtual drug screening, and elucidation of protein structure-function relationships in health and disease (McFedries *et al*., 2013, Maveyraud & Mourey, 2020). While Cryo-electron microscopy is well suited for high molecular weight proteins, X-ray crystallography remains the tool of choice for obtaining high-resolution structures of lower molecular weight proteins (Cooper *et al*., 2011), (Zhang & Liu, 2018). Since the introduction of X-ray crystallographic methods, approximately 10% of proteins attempted have resulted in diffraction-quality crystals (Dale *et al*., 2003). The protein crystallization process is generally laborious, time-consuming, and expensive because of this high failure rate (Dale *et al*., 2003). The recent advancement of Micro-electron diffraction of protein microcrystals also supports a need for improved protein crystallization methods (Shi *et al*., 2013). The continuous need for protein structure determination in various fields calls for a faster and more reliable technique for protein crystallization.

Protein crystallization chaperones can improve the success rate of protein crystallization. Currently-available protein crystallization chaperone strategies can be divided into four main types: 1) Noncovalent crystallization chaperones (e.g., Nanobodies and DARPINS) (Koide, 2009, Kovari *et al*., 1995) 2) host-guest lattices (e.g., EngBF and R1EN) (Ernst *et al*., 2019, Maita, 2018), 3) Fused monomeric crystallization chaperones (e.g., Glutathione S-transferase (GST), Maltose Binding Protein (MBP), thioredoxin (TRX), and green fluorescent protein (GFP)) (Uhlen *et al*., 1992, Malhotra, 2009), and 4) Polymer forming crystallization chaperones (e.g., TELSAM) (Nauli *et al*., 2007, Kim *et al*., 2001).

The “TELSAM” variant of the human Translocation ETS Leukemia (TEL) protein sterile alpha motif (SAM) domain is a polymer-forming crystallization chaperone that is soluble at pH > 8. However, when the pH decreases below 7, TELSAM forms a six-subunit-per-turn helical polymer that can enable fused target proteins to co-crystallize (Kim *et al*., 2001, Nauli *et al*., 2007, Nawarathnage *et al*., 2022). TELSAM reduces the entropic cost of crystal nucleation and growth by pre-freezing some degrees of freedom in the crystal lattice. Additionally, pre-ordering many thousands of copies of the target protein along the polymer results in substantial avidity to strengthen the subsequent target protein-target protein interactions that are formed when adjacent TELSAM polymers associate. As a result, TELSAM-target protein fusions readily form crystals. TELSAM is thus a compromise between host-guest lattices, which readily form crystals but as yet do not force the target protein to participate in crystal contacts, and fused monomeric crystallization chaperones, which force the target protein to participate in crystal contacts but are not guaranteed to form crystals (Nawarathnage *et al*., 2022).

There are two principal strategies to fuse TELSAM and a protein of interest. The first employs a flexible linker that allows different possible orientations of the fused protein in the final crystal lattice, which may facilitate crystallization. The higher flexibility may alternatively lead to a greater degree of disorder in the crystal, leading to poorer diffraction resolution and higher B-factors. The second strategy uses a rigid linker that imparts stability to the target protein orientation in an effort to improve crystal order. The reduced linker flexibility may however block access to target protein orientations optimal for crystallization, potentially reducing crystallization propensity or crystal quality (Nawarathnage *et al*., 2022). In our previous study (Nawarathnage *et al*., 2022), we discovered that TELSAM could increase the rate of crystallization, form crystals without direct contact between TELSAM polymers, and in some cases using only very minimal crystal contacts overall. Replicating our previous results with 1TEL-flex-vWa uncovered potential areas for improvement for TELSAM-mediated protein crystallization:

1. The linker sequence between TELSAM and the fusion protein defines the movement and flexibility of the fused protein in crystal. In the previous study, the 1TEL-flex-vWa linker consisted of a single alanine. In the resulting crystal lattice, this alanine adopted an α-helical conformation, extending the 1TEL C-terminal α-helix. The 2nd and 3rd amino acids of the vWa domain, Ala-Ala in this construct, preceded the first β-sheet of the vWa domain and adopted a more flexible conformation. No optimization of the 1TEL-vWa linker was carried out in this initial study.
2. A cleavable 10xHis-SUMO tag was included in the construct to increase protein solubility and to allow His tag removal before crystallization. The effects of SUMO tag cleavage and the SUMO protease enzyme activity on crystallization propensity or crystal quality were not tested in the previous study.
3. In the previous study, we discovered TELSAM can increase crystallization rate but did not produce crystals that diffracted to resolution better than 2.8 Å. Optimization of the crystallization or freezing conditions was not investigated.
4. Although we determined that 1TEL-AA-vWa could be crystallized at concentrations as low as 1 mg/mL, we did not determine the lower concentration limit for crystallization of this construct. Low-concentration protein crystallization using TELSAM is a promising avenue of investigation because it can potentially decrease batch sizes and the cost of protein crystallization, as well as enable the crystallization of poorly-soluble and difficult-to-produce proteins.

As reproducibility in protein crystallization is desirable for drug screening and structure-based drug design applications, we sought to understand and address these issues.

## 2. Results

### 2.1. 1TEL-AA-vWa (SUMO) does not reliably form diffracting crystals

We purified and crystallized five separate batches of 1TEL-flex-vWa (here termed 1TEL-AA-vWa (SUMO)) to determine the robustness and reproducibility of TELSAM-mediated crystallization. The 1st batch of 1TEL-AA-vWa generated large hexagonal crystals in 3 days in a single condition (100 mM Bis-Tris, pH 5.7, 3.0 M NaCl) with an average diffraction limit of 2.9 Å (2.8–3.0 Å), as previously-reported (Nawarathnage *et al*., 2022), while the 3rd and 4th batches developed thinner needle-shaped crystals which did not diffract. A 5th batch did not form crystals.

One 1TEL-AA-vWa (SUMO) batch resulted in crystals that featured target proteins in multiple orientations relative to their host 1TEL domains across chains in the asymmetric unit, in multiple orientations within a single chain, or were not located. The 2nd batch of 1TEL-AA-vWa (SUMO) generated a fan of rod-like crystals in 10 days in a single condition (100 mM sodium acetate, pH 4.6, 2.0 M sodium formate) (Figure 1A). Prior to mounting, this fan of crystals was soaked with a 10 mM solution of a drug candidate, PGM, in 10% DMSO (Doyaguez *et al*., 2019). One of these crystals diffracted to 3.3 Å resolution (Table 1). Molecular replacement with this 1TEL-AA-vWa (SUMO) crystal was carried out using PDB IDs: 2QAR (1TEL) and 1SHT (vWa) as search models, yielding a solution with 1TEL polymers along the 65 axes and two additional 1TEL polymers along the two 32 axes of the P65 unit cell (Figure 1B). The asymmetric unit thus contains one 1TEL domain (chain B) from a polymer lying on a 65 axis and two 1TEL domains (chains A and C) from a polymer lying along a 32 axis. The vWa domains corresponding to the chains A and B 1TEL domains were also located, but the density for the 65 axis vWa domain (chain B) was incomplete, suggesting that this chain B vWa domain may exist in at least two orientations relative to its host 1TEL domain (Figure 1C). The vWa domain corresponding to the chain C 1TEL domain (32 axis) in the asymmetric unit was not located, although there is ample space in the unit cell for it and ample unassigned electron density in the area where it is expected to be found. We began by modeling extended poly-alanine peptides into this unassigned electron density and later replacing these with UNX atoms modeled at the coordinates of the peptide Cα atoms, to approximate the area occupied by the various putative chain C vWa orientations (Figure 1D).

**Figure 1.**
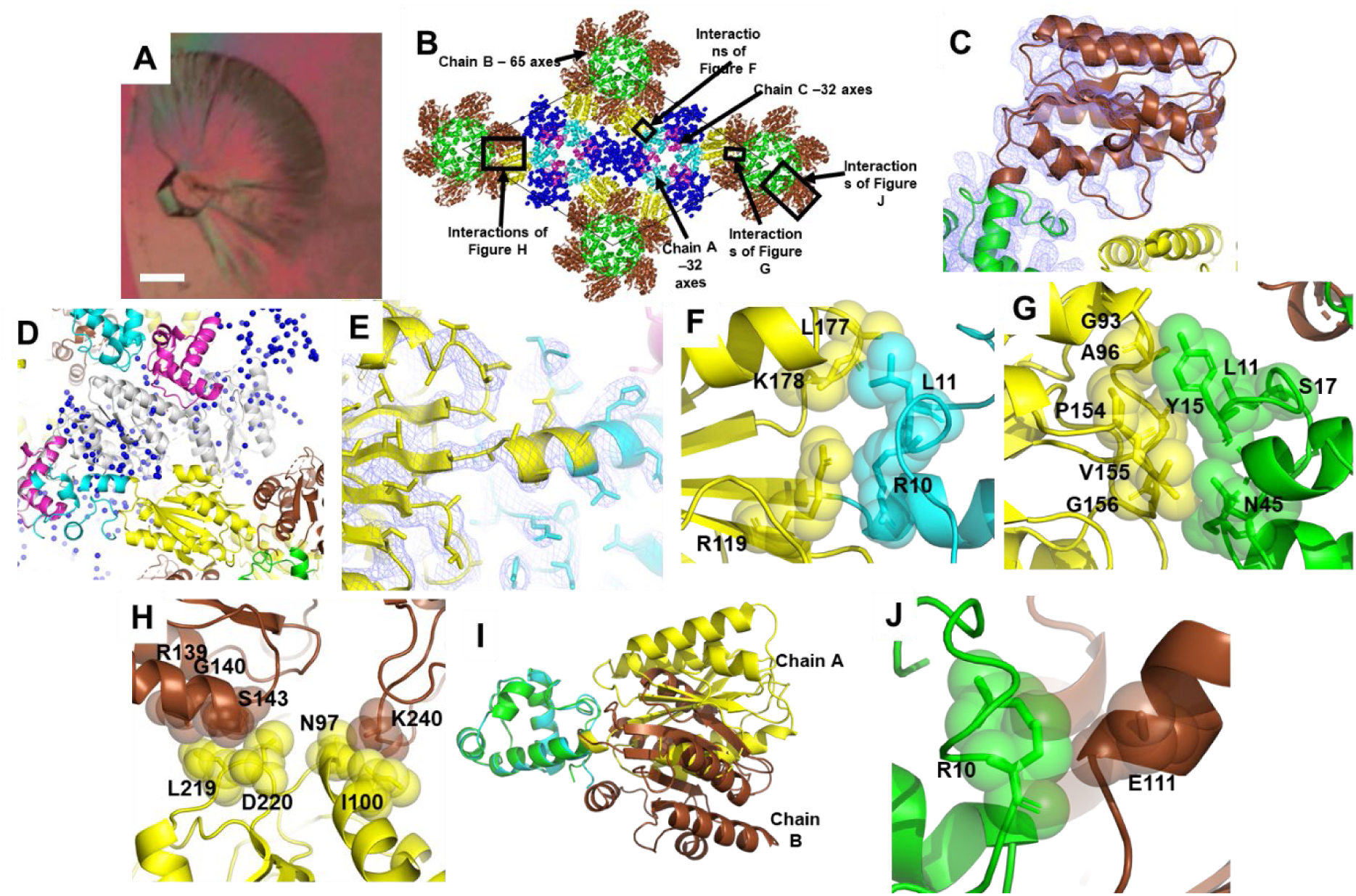
1TEL-AA-vWa (SUMO) crystal lattice. A. Batch 2 1TEL-AA-vWa (SUMO) crystal fan. The scale bar is 100 μm. B. Crystal lattice packing of the 1TEL-AA-vWa (SUMO) crystal. The TELSAM domains of chains A, B, and C are shown in Cyan, Green, and Magenta, respectively, while the vWa domains of chains A and B are shown in Yellow and Brown, respectively. The unit cell is delineated with a thin black line. C. Electron density (slate mesh) for the chain B vWa domain, contoured to 1.0 σ. The 1TEL (green) and vWa (brown) domains are shown in cartoon representation for reference. D. Detail of the region where the chain C vWa domain is expected to be found. A modelled orientation of the chain C vWa domain (white) is superimposed over the UNX atoms (blue spheres) used to delineate the unassigned electron density in this region. E. Detail of the linker between the chain A 1TEL (cyan) and vWa domains (yellow). The electron density is shown as a slate mesh and contoured to 1.0 σ. F. Detail of the interaction between the chain A vWa domain (yellow) and its host 1TEL domain (cyan). The proteins are shown in cartoon representation with selected amino acids shown as sticks and transparent spheres. G. As in F but of the interactions between the chain A vWa domain (yellow) and the chain B 1TEL domain (Green). H. As in F but of the interactions between the chain A vWa domain (yellow) and two chain B vWa domains (Brown). I. Superposition of chain A and chain B 1TEL-AA-vWa chains via their 1TEL domains. J. As in F but of the chain B vWa (brown) and its host 1TEL (green).

**Table 1.** Crystallization time, propensity, and diffraction quality of vWa constructs.

| Construct | Days to crystal appearance | Number of commercial conditions giving crystals | Number of commercial conditions screened | Diffraction limit of collected datasets(Å) | Fraction of non-ice reflections indexed (%) | Average maximum mosaicity used in data processing (°) | Final I/Sa of data processing | Unit cell solvent content (%) | Largest off origin Patterson peak (%) | Twin fraction (%) |
| --- | --- | --- | --- | --- | --- | --- | --- | --- | --- | --- |
| 1TEL-AA-vWa (SUMO)* | 3-10 | 9 (3%) | 288 | 2.9, n=3 (2.8–3.0) | 57 (43–67) | 0.47 (0.38–0.60) | 16 (15–17) | 59.0 | 5.06 | 0.00 |
| VWa alone* | 35-39 | 4 (1%) | 384 | 2.4, n=18 (1.9-2.9) | 56 (19 – 95) | 0.30 (0.20-0.45) | 38 (23 - 53) | 41.7 | 9.18 | 0.00 |
| 1TEL-AV-vWa (SUMO) | 3-10 | 4 (0.8%) | 480 | 2.9, n=8 (2.5 –3.5) | 35 (12 – 73) | 0.59 (0.23–1.1) | 20 (6–36) | 59.3 | 6.26 | 0.425 |
| 1TEL-TV-vWa (SUMO) | 3-10 | 4 (0.8%) | 480 | 2.7, n=6 (2.3 –3.0) | 49 (15 – 88) | 0.55 (0.18–1.1) | 19 (7–34) | 33.4 | 74.52 | 0.047 |
| 1TEL-AA-vWa | 7-10 | 2 (0.4%) | 480 | 2.6, n=4 (2.5 –2.8) | 67 (54 – 94) | 0.16 (0.07–0.21) | 19 (15–23) | 60.9 | 4.37 | 0.027 |
| 1TEL-TV-vWa | 2-7 | 4 (0.8%) | 480 | 2.0, n=16 (1.6 –2.6) | 77 (31 – 97) | 0.25 (0.12–0.40) | 25 (11–42) | 47.3 | 4.65 | 0.005 |

\* Data taken from (Nawarathnage *et al.*, 2022)

These observations suggest that the chain C vWa domain may exist in a variety of orientations relative to its host 1TEL polymer. Taken together, these data suggest that the vWa domains in this crystal may adopt at least six different orientations (one in chain A, at least two in chain B, and at least three in chain C). An alternative possibility is that interaction with the PGM drug molecule may have partially misfolded the chains B and C vWa domains. In this scenario, the chain B vWa domain may be partially resolved because it interacts with two copies of chain A vWa domains coming from two distinct 32-axis polymers.

We were able to refine this model to only relatively poor R-work/R-free values of 0.28/0.31, which diminishes confidence in the space group assignment and molecular replacement solution. Fortunately, TELSAM fusion crystal structures have two built-in sources of validation which bolster confidence in this space group assignment and solution: 1) The 1TEL monomers form the expected monomer–monomer interfaces and 6-fold left-handed helical polymer, 2) The N-termini of the vWa domains are within the expected distance of the C-termini of their host 1TEL monomers, and continuous electron density can be seen for the 1TEL–vWa linkers (Figure 1E). This is notable because molecular replacement placed the 1TEL and vWa domain search models as independent monomers and included no constraints to enforce the observed symmetry or contacts. No twinning or pseudosymmetry was detected in this crystal.

We note that the well-resolved chain A vWa domain (hosted by a 1TEL domain in the 32-axis polymer) adopts a novel binding mode to its host 1TEL polymer. For its evidently critical role in maintaining the integrity of the crystal, the chain A vWa domain makes surprisingly minimal contact with its host 1TEL domain, burying only 270 Å^2^ of solvent accessible surface area in a fairly solvated interface. Specifically, the vWa domain Arg 119, Leu 177, and Lys 178 make van der Waals contacts with the 1TEL Arg 10 and Leu 11 (Figure 1F).

The chain A vWa domain also makes direct contacts to the chain B 1TEL and vWa domains (in the 65-axis polymer), allowing the structural integrity of this crystal to be maintained (Figure 1B). These interactions are distinct from those observed in the previous Batch 1 structure of 1TEL-AA-vWa (SUMO) (PDB ID: 7N1O) (Nawarathnage *et al*., 2022). The chain A interface to the chain B 1TEL domain buries 252 Å^2^ of solvent-accessible surface area and is largely hydrophobic. Specifically, the side and main chains of the vWa domain Gly 93, Ala 96, Asn 97, Pro 154, Val 155, and Gly 156 form hydrophobic and van der Waals contacts to the side and main chains of the chain B 1TEL Leu 11, Tyr 15, Ser 17, and Asn 45 (Figure 1G). The chain A vWa domain interface with the chain B vWa domain buries 99 Å^2^ of solvent accessible surface area and is somewhat more polar. Specifically, the chain A vWa domain Leu 219 and Asp 220 side and main chains make hydrophobic and van der Waals interactions with the chain B vWa domain Arg 139, Gly 140, and Ser 143 (Figure 1H). This chain A vWa domain also forms contacts with a second chain B vWa domain from the same polymer as the first, burying 164 Å^2^ of solvent-accessible surface area. Specifically, the chain A vWa domain Asn 97 and Ile 100 make van der Waals interactions with Lys 240 from this second chain B vWa domain (Figure 1H).

The resolvable portions of the chain B vWa domain reveal that it adopts a binding mode to its host 1TEL polymer distinct from that of the chain A vWa domain but likely similar to the unresolved chain C vWa domain (Figure 1I). The chain B vWa domain makes only a glancing contact to its host 1TEL polymer involving van der Waals contacts between Glu 111 of the vWa domain and Arg 10 of the 1TEL domain and burying 120 Å^2^ of solvent accessible surface area (Figure 1J). This minimal contact may explain why the chain B vWa domain exists in more than one orientation relative to its host 1TEL domain.

As the three vWa domains in the asymmetric unit adopt at least six binding modes (clustered around two general binding modes) (Figure 1I), this Batch 2 1TEL-AA-vWa (SUMO) structure provides evidence that fused target proteins may be able to choose their binding modes to their host TELSAM polymers at the time of polymer-polymer association. Further evidence comes from the extremely weak interactions the chain A and B vWa domains make with their host 1TEL polymers. It is unlikely that these weak interactions would be made maintained prior to or in the absence of polymer-polymer association.

### 2.2. Decreasing the flexibility of the 1TEL--vWa linker improves diffraction resolution but does not improve crystal quality

In the batch 2 1TEL-AA-vWa (SUMO) crystal described in the previous paragraphs, we discovered the vWA domains in at least six different orientations among the three chains of the asymmetric unit, suggesting that the vWa does not readily find a single lowest-energy binding mode against its host 1TEL polymer during polymer-polymer association and that while failure to do so complicates structure solution, it does not necessarily abrogate the formation of diffracting crystals. We hypothesized that for TELSAM–target-protein fusions to form well-diffracting crystals with well-resolved target proteins, all of the fused target proteins must find the same binding mode against their host polymers so as to allow the TELSAM–target-protein polymers to reproducibly find a single binding mode to one another. We further hypothesized that modification of the 1TEL–vWa linker could limit the orientational flexibility of the vWa domain such that all of the vWa domains in a given crystal would more easily adopt the same binding mode against their host 1TEL polymers, thus being resolved in the resulting electron density and potentially increasing the diffraction resolution of the crystal.

The single Ala linker used in the previous 1TEL-AA-vWa (SUMO) structure (PDB ID: 7N1O) became part of the rigid C-terminal α-helix of the 1TEL domain. The 2nd and 3rd amino acids of the vWa domain (Ala-Ala in this construct) were seen in an extended conformation and did not pack well against the vWa amino acids around them, suggesting that they are the most flexible part of the 1TEL–vWa connection in this crystal, despite lying within the vWa rather than in the designed linker (Figure 2A) (Nawarathnage *et al*., 2022).

**Figure 2.**
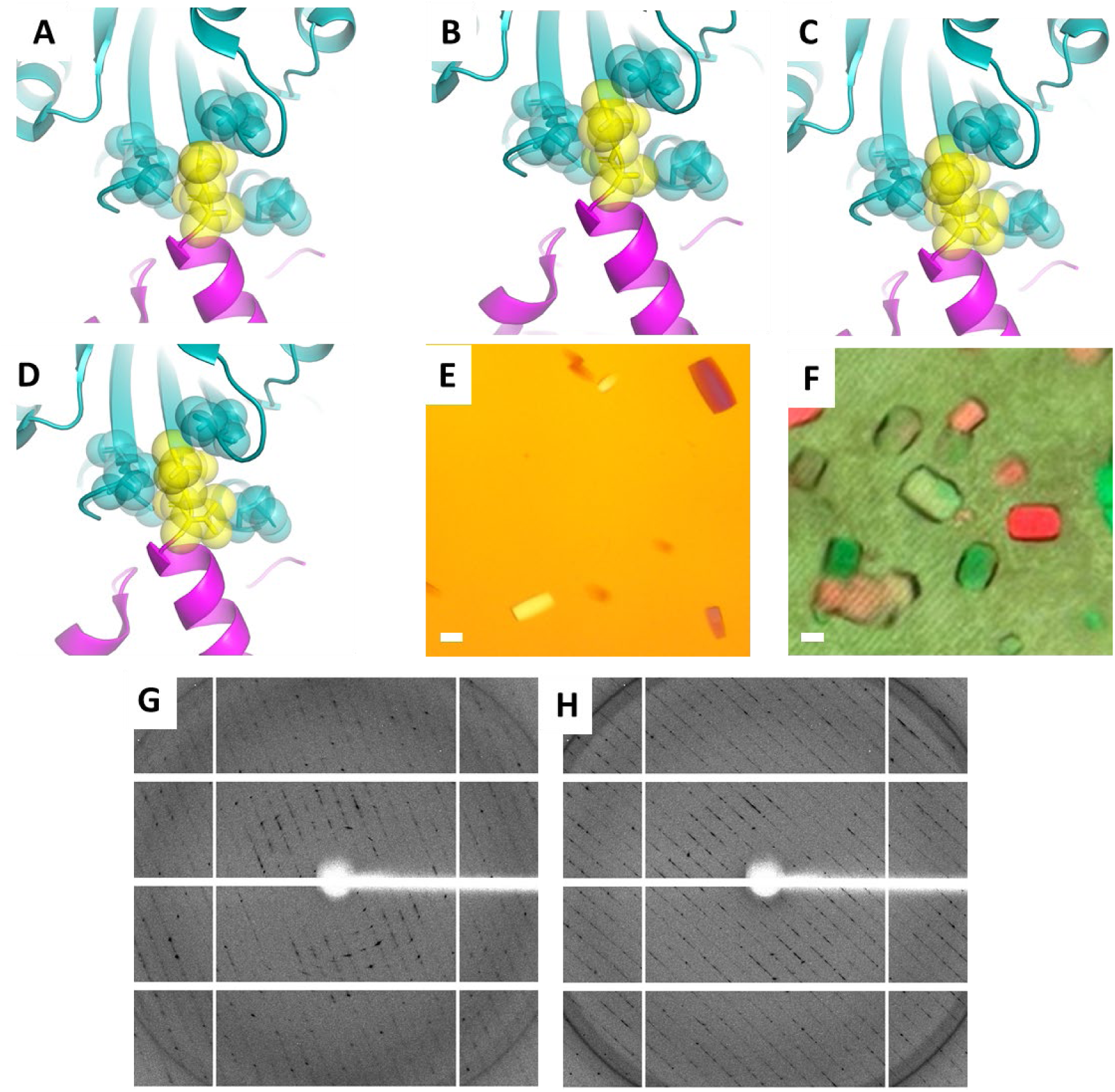
Results of adding bulk to the 1TEL–vWa linker. A. Linker detail of the 1TEL-AA-vWa structure (PDB ID: 7N1O) with the 1TEL in magenta, the Ala-Ala linker in yellow, and the vWa in cyan. Linker and surrounding amino acids are shown as sticks. B. As in A, but for the design model of 1TEL-AV-vWa. C. As in A but for the design model of 1TEL-TV-vWa. D. As in A but for the design model of 1TEL-TT-vWa. E Representative crystals of 1TEL-TV-vWa (SUMO). The scale bar is 100 µm. F. As in E but of 1TEL-AV-vWa (SUMO). G. Representative diffraction pattern of a 1TEL-TV-vWa crystal. H. As in G but of a 1TEL-AV-vWa crystal.

We substituted the Ala-Ala linker with either Ala-Val (AV), Thr-Val (TV), or Thr-Thr (TT). These substitutions packed better with amino acids around them and were intended to confer rigidity to the 1TEL--vWa connection (Figure 2B-D). We cloned the three different linker variants to create 1TEL-AV-vWa (SUMO), 1TEL-TV-vWa (SUMO), and 1TEL-TT-vWa (SUMO) constructs. We expressed, purified, and crystallized each construct at both 1 and 20 mg/mL with a range of commercially available and custom-made crystallization screens. We executed a single round of optimization of crystallization conditions but no optimization of cryoprotection or freezing conditions.

1TEL-AV-vWa (SUMO) and 1TEL-TV-vWa (SUMO) crystallized in 4 different conditions in 2-3 days, while 1TEL-TT-vWa (SUMO) did not crystallize (Figure 2E-F). We identified initial hits in commercial crystallization screens and from those hits developed an ammonium sulfate optimization screen. 1TEL-AV-vWa (SUMO) crystals diffracted to an average resolution of 2.9 Å (I/s ≥ 2.0, 2.5– 3.5 Å across 8 crystals). For these crystals we were able to index an average of 35% of non-ice reflections (12%–73%), with an average mosaicity of 0.59° (0.23–1.1°), and an average ISa of 20 (6– 36) (Diederichs, 2010) (Table 1). 1TEL-TV-vWa (SUMO) crystals diffracted to an average resolution of 2.7 Å (I/s ≥ 2.0, 2.3– 3.0 Å across 6 crystals). For these crystals, we were able to index and average of 49% of non-ice reflections (15–88%), with an average mosaicity of 0.55° (0.18–1.1°), and an average ISa of 19 (7–34).

By comparison, the Batch 1 1TEL-AA-vWa (SUMO) crystals gave an average resolution of 2.9 A (2.8–3.0 A across 3 crystals), indexed 57% of non-ice reflections (43-67%), exhibited an average mosaicity of 0.47° (0.38--0.60°) and had an average ISa of 16 (15-17) (Nawarathnage *et al*., 2022). These results suggest that the Thr-Val linker performed slightly better than the Ala-Ala linker, which in turn performed on par with the Ala-Val linker, while the Thr-Thr linker abrogated crystal formation.

Some 1TEL-vWa crystals feature flipped TELSAM-target polymers. The 1TEL-TV-vWa (SUMO) crystal and dataset selected for structure solution had a larger P65 unit cell (a = b = 164.156 Å, c= 54.409 Å) than other 1TEL-TV-vWa (SUMO) and 1TEL-AV-vWa (SUMO) crystals, similar to the unit cell parameters of the Batch 2 1TEL-AA-vWa (SUMO) crystal (a = b = 162.557 Å, c = 56.877 A) (Figure 1A). This dataset also exhibited significant translational non-crystallographic symmetry (tNCS) (the largest off-origin peak in the Patterson map was 74.52% of the height of the origin peak). Molecular replacement with this dataset was carried out as before, once again yielding a solution with 1TEL polymers along the 65 axes and two additional 1TEL polymers along the two 32 axes of the unit cell. In this case we did not see multiple orientations of the vWa domains relative to their host 1TEL polymers (Figure 3A, B). For ease of explanation, we term these three chains Chain A (1TEL polymers on the 32 axes of the unit cell whose vWa domains interact with vWa domains from 1TEL polymers (chain B) on the 65 axes), Chain B (1TEL polymers on the 65 axes of the unit cell whose vWa domains interact with vWa domains from 1TEL polymers (chain A) on the 32 axes), and Chain C (1TEL polymers on the 32 axes whose vWa domains interact with their symmetry equivalent 32-axis vWa domains. Our initial molecular replacement solution featured parallel polymers that all ran in the same N→C direction, but we were not able to refine it to acceptable R-factors (Rwork/Rfree = 0.37/0.38). We next tried reindexing this dataset into a smaller unit cell with 1TEL polymers only on the 65 axes. This likewise gave only poor Rfactors (Rwork/Rfree = 0.33/0.35). We attempted twin refinement in PhenixRefine using the twin operator h,-h-k,-l, (the two-fold axis parallel to **a**) for each unit cell size, but this likewise yielded poor R-factors. The fact that twin refinement failed to improve the fit to the data suggests that polymers with different N→C directions are uniformly distributed over the crystal rather than aggregated in separate twin domains.

**Figure 3.**
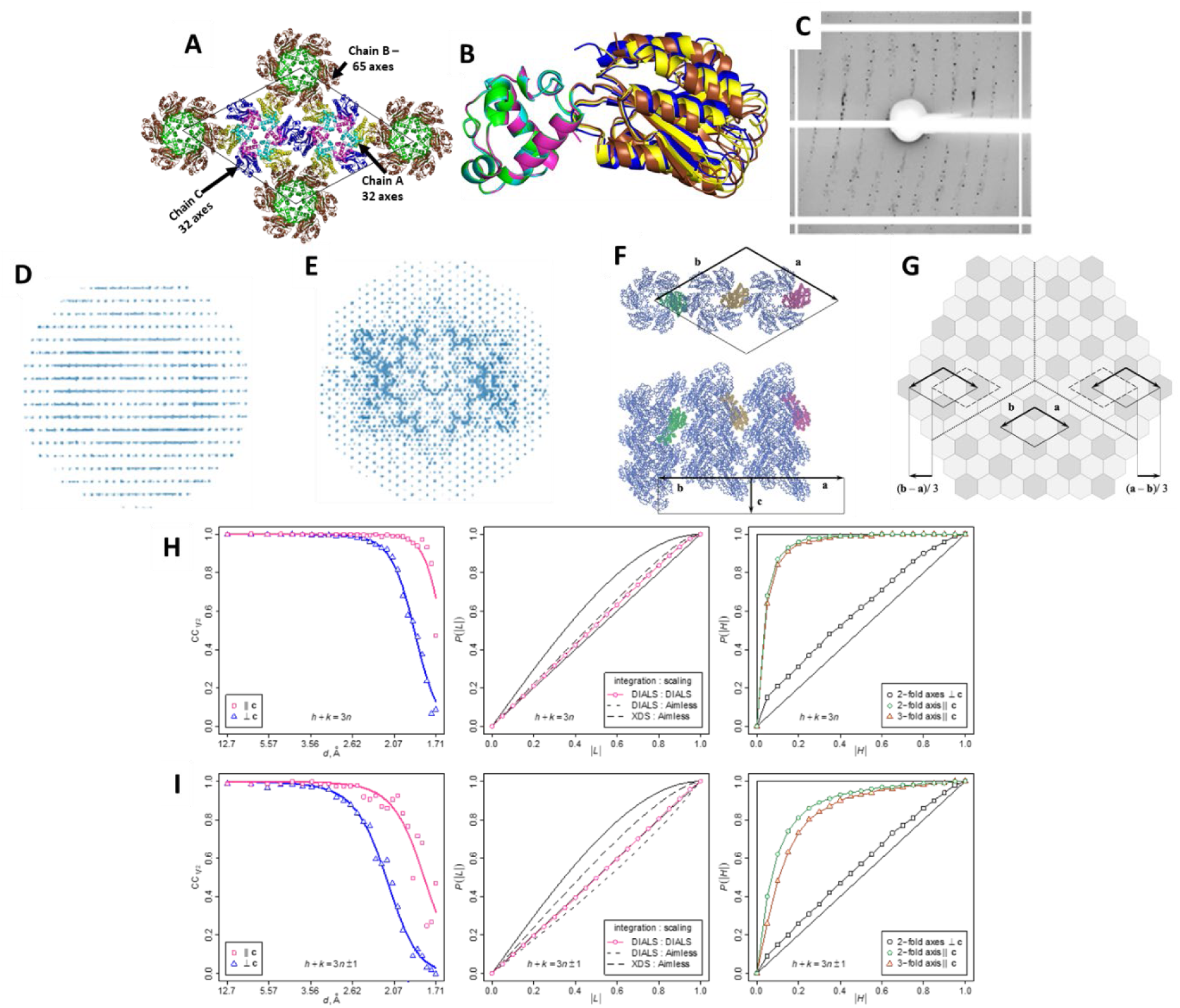
Possible models of crystal disorder. A. Lattice contacts in the unit cell, with proteins shown in cartoon representation. Chain A (32-axis) is colored cyan (1TEL) and yellow (vWa). Chain B (65-axis) is colored green (1TEL) and brown (vWa). Chain C (32-axis) is colored magenta (1TEL) and purple (vWa). The unit cell is shown as a thin black line. B. As in A, but showing the chain A, B, and C 1TEL-TV-vWa domains superimposed through their 1TEL domains. C. Representative diffraction pattern of the 1TEL-TV-vWa (SUMO) crystal, showing diffuse scattering between the strong reflections. D. Projection of the reciprocal space viewed perpendicular to **c**. E. as in D but viewed along **c**. F. Packing ambiguity of adjacent helical polymers. G. Possible model of crystal architecture. The bright and dark hexagons represent 32 (chains A and C) and 65 (B chain) axis polymers, respectively. On the large scale, the difference between 32 and 65 polymers is that they are related by two-fold rotation around an axis in the plane of the figure as shown in F, and on a smaller scale, the relative orientation of the 1TEL and vWa protein domains differ by 6.8° (chains A and B), 5.8° (A and C), 11.7° (C and B). I-J. Comparison of intensity statistics of reflection subsets. Top row: reflections h - k = 3n. Bottom row: reflections h - k = 3n +/- 1. Left column: CC1/2 show considerable differences in the useful resolution range of the two subsets. Center column: The results of the L-test for twinning strongly depend on data processing. Right column: The results of H-tests are consistent between the two subsets of reflections and indicate no twinning relative to the two-fold axis perpendicular to **c**. Thin solid black lines in the L-test plots represent theoretical curves for twinned (top) and non-twinned data (bottom). Thin black lines in the H-test plots represent the theoretical curve for the twinning operation or crystallographic symmetry operation (top line, step function) and the lattice symmetry operation unrelated to crystal symmetry or twinning (bottom line).

We returned to the 1TEL-TV-vWa (SUMO) diffraction dataset to identify the cause of our poor refinement statistics and of the signs of an extreme tNCS (non-origin Patterson peak with a height 74.52% of the origin peak). The diffraction data for this 1TEL-TV-vWa (SUMO) crystal featured diffuse scattering between two out of every three reflections (Figure 3C-E). We hypothesized that this crystal is formed by small domains with symmetry P65, packed with an offset of (a-b)/3 relative to their neighbors.

With this model, we attribute the diffuse patterns in the diffraction images (Figure 3C-E) to broken global periodicity in the crystal. In published structures with monoclinic ordered domains, streaky diffraction spots were observed, with streaks extending in the direction of broken crystal periodicity, and with rows of streaky reflections alternating with rows of well-defined reflections in one of two other directions. Accordingly, the integrated intensities of reflections were modulated in the latter direction, with the period of modulation being non-commensurate with the reciprocal lattice. The diffraction images of the 1TEL-TV-vWa (SUMO) crystal show two-dimensional diffuse patterns in the planes perpendicular to **c\*** (Figure 3D). The streaks connect weak reflections h-k = 3n+/-1 forming hexagonal shapes around well-defined strong reflections h-k = 3n. Here, the modulation of intensities is consistent with a hexagonal lattice and the modulation coefficients take only two values, one applies to reflections h-k = 3n and the other to the reflections h-k = 3n+/-1. The high ratio of the two coefficients of about 7 (see Materials and Methods) explains the high values of the R-factors in refinements using all of the data without correction.

The structure and P65 symmetry of the ordered domains were determined by molecular replacement using а subset of strong reflections and confirmed later by molecular replacement against a complete but demodulated dataset expanded to space group P1. The domains are formed by helical polymers of 1T-TV-vWa (SUMO) molecules, with the 65 axes polymers coinciding with the crystallographic unit cell axes. TELSAM helical polymers are present in two orientations; one is adopted by the polymers lying on the 32 crystallographic axes (further referred to as 32-axes polymers) and the other by the polymers lying on the 65 axes (65-axes polymers) (Figure 3F). This explains the high R-factors in the refinement in the small cell, wherein the model corresponds to a putative crystal packing with all polymer helices in the same orientation.

The 32-axis polymers form a continuous interconnected framework, whereas the 65 helices make no contact with each other. In the model in Figure 3G, these requirements are satisfied in each of the ordered domains and in the interfaces between them. Were it not for this constraint, there would be nothing to prevent some of the adjacent domains from twinning, However, the H-test for two-fold axes perpendicular to **c** indicates the opposite (Figure 3I, J, right panels). High values of R-merge for these symmetry operations, above 0.55 for resolution cutoffs 2.2 and 2.9 Å, are in agreement with the H-test. We thus favor the model in Figure 3G with the direct contacts between 65-axes polymers prohibited.

The translation (a-b)/3 is a tNCS operation in the 1TEL-TV-vWa (SUMO) structure because it relates some 32-polymers with some other 32-polymers, but it is not a pseudosymmetry operation, because it places 65 polymers in the positions of the 32 polymers in an incorrect orientation. Therefore, the structure cannot be approximated by a structure with a three times smaller unit cell, and correction of intensities was critical for structure determination. However, it is easy to imagine a structure with pseudosymmetry (a-b)/3 and also with ordered domains offset by this operation. The structure solution in the small cell would then be possible and the electron density maps would represent an average density from the three molecules related by pseudosymmetry. The weak reflections could still be observed but refinement of a structure with correct unit cell parameters to reasonable R-factors would be impossible (without correction of intensities). It is likely that difficulties with refinement in e.g. (Ostrowski *et al*., 2015) and unexplained weak reflection observed in e.g. (Li *et al*., 2016) can be explained by crystal defects similar to those in the 1TEL-TV-vWa (SUMO) crystal. The corrected set of structure amplitudes is deposited with the PDB along with the scaled unmerged data without correction. The ISa, R-work and R-free for the subset of reflections h - k = 3n known to be least affected by partial crystal disorder are provided in Table 2 for reference.

**Table 2.** Data collection and refinement statistics.

| Construct | Batch 2<br>1TEL-AA- | 1TEL-TV-vWa | 1TEL-TV-vWa | 1TEL-AA-vWa | 1TEL-TV-vWa | 1TEL-<br>helix- | 1TEL-<br>G-Ack1- | vWa<br>alone |
| --- | --- | --- | --- | --- | --- | --- | --- | --- |

|  | vWa<br>(SUMO)<br>PDB ID:<br>8FZV | (SUMO)<br>PDB ID:<br>8FZU | (SUMO)<br>(1/3 of<br>data) | PDB ID:<br>8FT6 | PDB ID:<br>8FT8 | TNK1-UBA<br>PDB ID:<br>8E1F | UBA<br>PDB ID:<br>8FZ3 | PDB ID:<br>8FZ4 |
| --- | --- | --- | --- | --- | --- | --- | --- | --- |
| **Wavelength (Å)** | 0.979460 | 1.00000 | 1.00000 | 1.85321 | 0.979460 | 0.97946<br>0 | 0.97946<br>0 | 0.97946<br>0 |
| **Resolution range**<br>**(Å)** | 70.39 - 3.292<br>(3.409 -<br>3.292) | 71.06 -<br>1.9 (1.968<br>- 1.9) | 82.04 -<br>1.9 | 43.19 -<br>2.62<br>(2.714 -<br>2.62) | 19.87 -<br>1.6<br>(1.658 -<br>1.6) | 33.02 -<br>2.16<br>(2.237 -<br>2.16) | 44.84 -<br>2.784<br>(2.883 -<br>2.784) | 39.08 -<br>2.19<br>(2.268 -<br>2.19) |
| **Space group** | P 65 | P 65 | P 65 | P 65 | P 65 | P 65 | P 65 | C 1 2 1 |
| **Unit cell** | 162.557<br>162.557<br>56.877 90 90<br>120 | 164.096<br>164.096<br>54.4009 90<br>90 120 | 164.096<br>164.096<br>54.4009<br>90 90 120 | 99.745<br>99.745<br>50.135 90<br>90 120 | 100.882<br>100.882<br>49.754 90<br>90 120 | 61.954<br>61.954<br>167.574<br>90 90<br>120 | 61.506<br>61.506<br>166.139<br>90 90<br>120 | 78.244<br>88.969<br>59.779<br>90<br>128.127<br>90 |
| **Total reflections** | 101038<br>(10665) | 668275<br>(67759) | 308068<br>(15372) | 296744<br>(28503) | 724353<br>(74091) | 387440<br>(37702) | 178539<br>(18870) | 113715<br>(11309) |
| **Unique**<br>**reflections** | 12820 (515) | 66252<br>(2186) | 30824<br>(1592) | 8712<br>(861) | 38098<br>(3766) | 19209<br>(1954) | 8929<br>(898) | 16403<br>(1620) |
| **Multiplicity** | 7.9 (8.1) | 10.1 (10.3) | 10.0 (9.7) | 34.1<br>(33.0) | 19.0<br>(19.7) | 20.2<br>(19.3) | 20.0<br>(21.0) | 6.9 (7.0) |
| **Completeness**<br>**(%)** | 90.65 (39.00) | 90.48<br>(33.33) | 99.9<br>(98.5) | 99.91<br>(100.00) | 99.56<br>(99.13) | 98.47<br>(100.00) | 99.84<br>(100.00) | 98.56<br>(98.96) |
| **Mean I/σ(I)** | 6.02 (1.12) | 4.67 (0.41) | 8.8 (0.3) | 27.46<br>(2.95) | 25.28<br>(2.54) | 17.04<br>(2.64) | 20.00<br>(2.60) | 10.80<br>(2.38) |
| **Wilson B-factor**<br>**(Å²)** | 118.66 | 24.16 | 29.03 | 62.84 | 27.12 | 33.66 | 63.22 | 35.43 |
| **R-merge** | 0.3371<br>(3.593) | 0.1787<br>(2.574) | 0.120<br>(3.669) | 0.1169<br>(1.255) | 0.05639<br>(0.9895) | 0.1397<br>(1.488) | 0.1248<br>(1.48) | 0.08972<br>(0.8622) |
| **R-meas** | 0.3611 | 0.1884 | 0.126 | 0.1187 | 0.05799 | 0.1433 | 0.1281 | 0.09702 |
|  | (3.838) | (2.709) | (3.879) | (1.275) | (1.015) | (1.528) | (1.516) | (0.9318) |
| **R-pim** | 0.1272<br>(1.331) | 0.05901<br>(0.8398) | 0.040<br>(1.241) | 0.0202<br>(0.2203) | 0.01339<br>(0.2253) | 0.0318<br>(0.3457) | 0.02867<br>(0.3301) | 0.0365<br>(0.3496) |
| **CC1/2** | 0.99 (0.283) | 0.998<br>(0.607) | 0.999<br>(0.092) | 1 (0.853) | 0.999<br>(0.939) | 0.999<br>(0.829) | 1 (0.864) | 0.999<br>(0.849) |
| **CC\*** | 0.998 (0.664) | 1 (0.869) | 1 (0.410) | 1 (0.96) | 1 (0.984) | 1 (0.952) | 1 (0.963) | 1 (0.958) |
| **Reflections used**<br>**in refinement** | 12056 (516) | 59954<br>(2186) | 22075<br>(2193) | 8708<br>(863) | 38073<br>(3762) | 19184<br>(1951) | 8918<br>(898) | 16390<br>(1620) |
| **Reflections used**<br>**for R-free** | 595 (24) | 3116 (125) | 1092(54) | 400 (45) | 1951<br>(204) | 930 (81) | 467 (38) | 782 (77) |
| **R-work** | 0.2823<br>(0.3762) | 0.2375<br>(0.3108) | 0.2347<br>(0.2983) | 0.2021<br>(0.2901) | 0.1820<br>(0.2585) | 0.1975<br>(0.2622) | 0.2285<br>(0.3648) | 0.2172<br>(0.3178) |
| **R-free** | 0.3145<br>(0.3550) | 0.2663<br>(0.3346) | 0.2827<br>(0.3800) | 0.2298<br>(0.2825) | 0.2042<br>(0.2878) | 0.2288<br>(0.3316) | 0.2662<br>(0.3348) | 0.2552<br>(0.3522) |
| **CC (work)** | 0.468 (0.018) | 0.827<br>(0.753) | 0.799(0.6<br>58) | 0.955<br>(0.753) | 0.963<br>(0.923) | 0.962<br>(0.850) | 0.971<br>(0.726) | 0.957<br>(0.873) |
| **CC (free)** | 0.471 (0.132) | 0.781<br>(0.691) | 0.673(0.5<br>19) | 0.919<br>(0.769) | 0.962<br>(0.899) | 0.930<br>(0.813) | 0.949<br>(0.585) | 0.935<br>(0.777) |
| **Number of non-**<br>**hydrogen atoms** | 4021 | 6714 | 5187 | 1920 | 2119 | 2788 | 2484 | 2606 |
| **macromolecules** | 3926 | 5881 | 9828 | 1868 | 1960 | 2572 | 2446 | 2505 |
| **ligands** | 91 | 6 | 0 | 20 | 22 | 18 | 7 | 4 |
| **solvent** | 4 | 827 | 22 | 32 | 144 | 198 | 31 | 97 |
| **Protein residues** | 567 | 761 | 716 | 255 | 255 | 318 | 312 | 354 |
| **RMS(bonds)** | 0.002 | 0.004 | 0.0030 | 0.004 | 0.006 | 0.012 | 0.008 | 0.002 |
| **RMS(angles)** | 0.36 | 0.66 | 0.59 | 0.57 | 0.76 | 1.09 | 0.84 | 0.47 |
| **Ramachandran**<br>**favored (%)** | 92.74 | 96.56 | 95.38 % | 96.81 | 97.21 | 99.03 | 98.36 | 96.86 |
| **Ramachandran**<br>**allowed (%)** | 6.53 | 3.44 | 4.62 % | 3.19 | 2.79 | 0.97 | 1.64 | 3.14 |
| **Ramachandran outliers (%)** | 0.73 | 0.00 | 0.00 % | 0.00 | 0.00 | 0.00 | 0.00 | 0.00 |
| **Rotamer outliers (%)** | 0.32 | 0.17 | 0.22 % | 0.00 | 0.00 | 0.00 | 0.00 | 0.00 |
| **Clashscore** | 4.61 | 5.77 | 1.22 | 5.83 | 2.57 | 3.02 | 10.22 | 1.88 |
| **Average B-factor (Å<sup>2</sup>)** | 121.67 | 29.15 | 35.13 | 65.07 | 41.57 | 43.93 | 68.27 | 52.33 |
| **macromolecules** | 121.54 | 28.53 | 38.39 | 64.95 | 40.93 | 43.52 | 68.40 | 52.38 |
| **ligands** | 128.82 | 48.85 | N/A | 82.66 | 69.49 | 64.26 | 93.19 | 56.14 |
| **solvent** | 82.01 | 33.39 | 28.47 | 60.94 | 47.30 | 47.40 | 52.59 | 50.68 |
| **Number of TLS groups** | 10 | 10 | 16 | 2 | 0 | 21 | 18 | 12 |

We executed molecular replacement using both a version of the dataset that included only the strongest 1/3 of the reflections and a version that included all reflections, with the strongest 1/3 of reflections having a different scale factor then than the remaining 2/3 of reflections, both cut back to 1.9 Å resolution. Molecular replacement was executed as before for both versions of the data and produced essentially the same solution in each case, placing 1TEL polymers on both the 65 and 32 axes of the unit cell as before. As expected, the 1TEL polymers on the 65 axes (chain B) of the unit cell ran in the opposite N→C direction from the polymers on the 32 axis, consistent with our model of the overall crystal architecture (Figure 3F, G). The vWa domains in this model were all fully resolved (in contrast to the Batch 2 1TEL-AA-vWa (SUMO) structure) and all adopted the same binding mode against their host 1TEL polymers, regardless of the polymer orientation (Figure 3B). This binding mode was distinct from any of the vWa binding modes seen in either the Batch 1 (PDB ID: 7N1O) or Batch 2 (PDB ID: 8FZU) 1TEL-AA-vWa (SUMO) structures (Figure 7E).

We were able to refine the model against the 1/3 dataset to Rwork/Rfree 0.23/0.28 and against the complete dataset to 0.24/0.27 (Table 2). The vWa domains (using chain A as an example) adopt a binding mode against their host 1TEL domains that buries 326 Å^2^ of solvent-accessible surface area (average of both sides of the interface). Specifically, the side chains of Leu 177, Gly 176, Ala 174 and Thr 179 of the vWa domain interact with the side chains of Gln 78, Thr 81 and Val 82 of the 1TEL domain (Figure 4A). The chain A vWa makes glancing contacts to the chain C vWa domains on either side of it within its host polymer, as well as to the chain A 1TEL domain one turn of its host polymer above it. This last interface buries 188 Å2 of solvent-accessible surface area and is mostly polar and highly solvated. Specifically, this interface involves polar and van der Waals contacts between the vWa Asp 148 and Arg 151 and the 1TEL Arg 18, Asp 19, Ala 22, and Asn 40 (Figure 4B). The chain A vWa domain makes no contacts to the 1TEL domains of adjacent polymers, similar to the Batch 1 1TEL-AA-vWa (SUMO) structure (PDB ID: 7N1O), but in contrast to the Batch 2 1TEL-AA-vWa (SUMO) structure.

**Figure 4.**
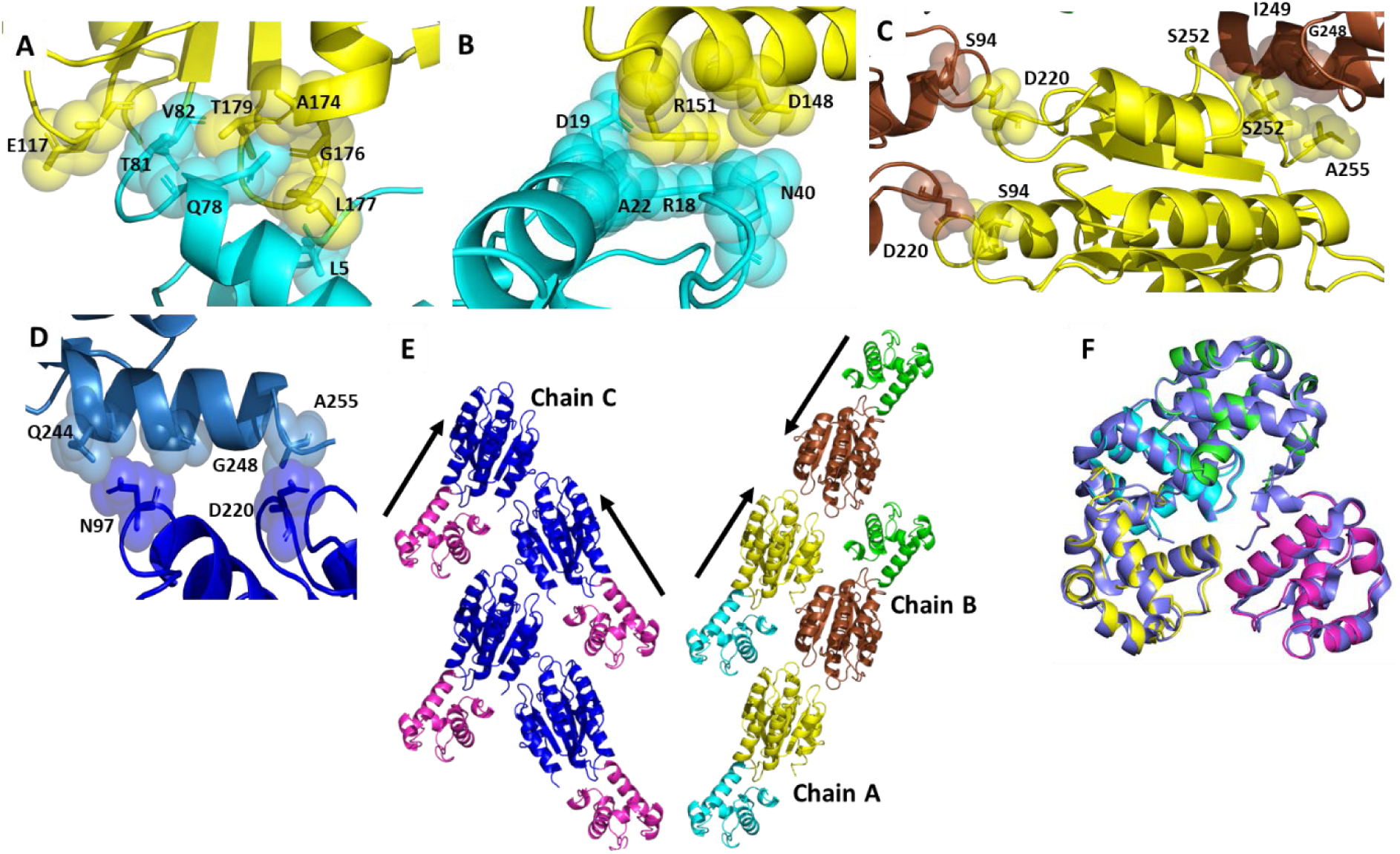
1TEL-TV-vWa (SUMO) molecular details. A. Detail of the interaction between the chain A vWa domain (yellow) and its host 1TEL domain (cyan). The protein is shown in cartoon representation with selected side chains shown as sticks and transparent spheres. B. As in A, but for the interaction between the chain A vWa and a 1TEL domain from the next turn of the host polymer. C. As in A, but for the interaction between the chain A vWa domain (yellow) and two chain B vWa domains (brown). D. As in A, but for the interaction between the chain C vWa domain (light blue) and an adjacent, symmetry-related chain C vWa domain (dark blue). E. Comparison of the antiparallel polymer-polymer interaction between chain A (cyan and yellow) and chain B (green and brown) and the parallel polymer-polymer interaction between 2 copies of chain C. F. Superposition of the unit cell contents of the 1TEL-helix-TNK1-UBA structure (slate) and the 1TEL-G-Ack1 structure (green, cyan, magenta, and yellow).

1TEL-TV-vWa (SUMO) provides an example where all target proteins adopt the same binding mode against their host polymers, but adjacent polymers adopt one of two binding modes to one another. This observation suggests that while the vWa binding mode to its host 1TEL polymer can be selected at the time of polymer-polymer association, at least in the case of 1TEL-TV-vWa (SUMO) there exists a sufficiently low-energy binding mode such that all of the vWa domains reliably select the same binding mode, possibly even some time prior to polymer-polymer association.

To investigate the cause of TELSAM polymer flipping in this crystal, we analyzed the contacts made between parallel polymers and those made between antiparallel polymers. The 32-axis chain A vWa domains make C2 symmetric “head-to-head” interactions with 65-axis chain B vWa domains, burying 341 Å^2^ of solvent-accessible surface area. This interface comprises a mixture of hydrogen bonds, hydrophobic interactions, and van der Waals interactions. Specifically, Ser 94, Asn 97, Asn 98, Glu 101, Leu 219, Asp 220, and Phe 221 on the chain A vWa domain contact the very same set of residues on the chain B vWa domain. Numerous well-ordered water molecules are also seen both within and around the periphery of this interface (Figure 4C).

Along the length of the TELSAM polymers, the 32-axis chain C vWa domains adopt a zipper-like arrangement, with each vWa domain contacting two other chain C vWa domains from an adjacent 32-axis polymer. Each vWa domain-vWa domain interaction buries 209 Å^2^ of solvent-accessible surface area and consists mostly of hydrophobic and van der Waals contacts between the side and main chains of Gln 244, Gly 248, Asn 251, Ser 252, and Ala 255 of one vWa domain and Asn 97, Asn 98, and Leu 219 of the second vWa domain. Numerous well-ordered water molecules and a sulfate ion are also seen around the periphery of this interface (Figure 4D). When the solvent content of the Chain A vWa to Chain B vWa interface is considered, we propose that the antiparallel chain A to chain B interface is qualitatively similar in estimated binding energy to the more hydrophobic parallel chain C to chain C interface, enabling the observed polymer flipping phenomenon (Figure 3F, 4E). Similar pathologies were also seen in other 1TEL-TV-vWa (SUMO) and 1TEL-AV-vWa (SUMO) crystals (Table 1).

### 2.3. Replacing the cleavable 10xHis-SUMO tag with a non-cleavable 10xHis tag significantly improved diffraction resolution and correlates with an absence of TELSAM polymer flipping

Decreasing the flexibility of the 1TEL–vWa linker moderately improved the diffraction resolution but failed to prevent other pathologies, such as twinning, tNCS, and TESAM polymer flipping. We have observed that trace proteases can cleave off portions of target proteins or even entire target proteins and that this can have profound effects on TELSAM polymerization and crystallization dynamics and architecture, possibly due to target protein cleavage products extending from the 1TEL C-terminus making transient interactions with other 1TEL fusion molecules and/or interfering with canonical TELSAM polymer formation.

For example, we attempted to crystallize a 1TEL-helix-TNK1-UBA (SUMO) construct. Crystals appeared in only a single condition (200 mM MgSO4, pH 5.9, 20% w/v Polyethylene Glycol 3,350) and diffracted to an average resolution of 2.08 Å (1.94-2.30 Å resolution across 3 crystals), with an average fraction of non-ice reflections indexed of 49% (43-56%), an average ISa of 14 (12-17), and an average mosaicity of 0.40° (0.26-0.56°). Molecular replacement using PDB IDs 2QAR and 7TCY as search models successfully located 24 copies of 1TEL in the P65 unit cell but not the UBA domain. In addition, the 1TEL domains did not form the expected left-handed helical polymer (Figure 4F).

In a second example, we attempted to crystallize a 1TEL-Gly-Ack1-UBA domain construct. In this case, the SUMO domain was omitted and a non-cleavable 10xHis tag was appended directly to the 1TEL N-terminus. Large crystals appeared in a single condition (1.4 M NaH2PO4/KHPO4, pH 8.2) and diffracted to an average resolution of 2.41 Å (2.08 and 2.78 Å resolution across 2 crystals), with an average fraction of non-ice reflections indexed of 44% (42 and 48%), an average ISa of 17 (11 and 22), and an average mosaicity of 0.56° (0.48° and 0.64°). Molecular replacement using PDB IDs 2QAR a homology model of the Ack1 UBA domain as search models again located 24 copies of 1TEL in the unit cell but not the UBA domain. Likewise, the 1TEL domains failed to form the expected left-handed helical polymer and instead adopted essentially identical crystal packing to the 1TEL-helix-TNK1-UBA construct described above (Figure 4F). This is striking because the SUMO tag and the tag cleavage step were omitted from the purification of this second construct. A PAGE gel of washed crystals revealed that this construct was intact; not cleaved by trace proteases. Our failure to locate the Ack1 UBA domain in the unit cell suggests that this domain might be unfolded and present in the unit cell as an extended disordered peptide.

The SUMO domain was originally included in our constructs to increase the solubility of the fusion protein and to allow the 10xHis tag to be removed using a thermophilic SUMO protease (Lau *et al*., 2018). The SUMO protease was expected to be more specific than other proteases commonly used for purification tag removal. In addition, the His-tagged protease is removed following tag cleavage by subsequently passing the cleavage reaction over a nickel-NTA column. Cleavage of target proteins from TELSAM may be due to nonspecific activity of trace SUMO protease not captured by the nickel-NTA step or by trace amounts of one or more other proteases present in the E. coli expression strain. Low levels of protease activity can result in 1TEL domains lacking their fused target proteins and peptide cleavage products being present in the crystallization drop, either of which might be responsible for the pathologies observed in the 1TEL–vWa (SUMO) constructs.

To avoid the possibility of nonspecific digestion of the target protein by trace proteases, we remade two genes, 1TEL-AA-vWa and 1TEL-TV-vWa, without the SUMO tag. We fused the 10xHis tag directly to the N-terminus of 1TEL and retained the tag through purification and into the crystal trays. We expressed, purified, and crystallized both proteins at 1 and 20 mg/mL with different commercially available and custom-made crystallization screens. 1TEL-TV-vWa formed crystals in 2-4 days, while 1TEL-AA-vWa took 7 days to crystallize. Some 1TEL-AA-vWa crystals were soaked with iodine before harvesting (Miyatake *et al*., 2006), while 1TEL-TV-vWa crystals were harvested without iodine treatment. Omission of the SUMO tag correlated with an improved diffraction resolution of 1TEL-AA-vWa and 1TEL-TV-vWa crystals by 0.4 Å and 0.7 Å, respectively. No twinning, pseudo-translation, or TELSAM polymer flipping was detected in diffraction datasets of these SUMO tag-free constructs.

1TEL-AA-vWa formed around 50 crystals averaging 300 µm in length (Figure 5A). The average resolution of these 1TEL-AA-vWa crystals was 2.6 Å (2.5--2.8 Å across 4 crystals) which was somewhat better than that of 1TEL-AA-vWa (SUMO), which was 2.9 Å (2.8-3.0 Å across 3 crystals) (Figure 5B, Table 1). Crystals of 1TEL-AA-vWa indexed an average of 67% of non-ice reflections (54-94%), exhibited an average mosaicity of 0.16° (0.07–0.21°) and an average ISa of 19 (15-23). Molecular replacement yielded a solution with 1TEL polymers along the 6-fold axes of the P65 unit cell with unit cell dimensions of a = b = 99.7 Å, c = 50.1 Å (Figure 5D). The vWa domains of 1TEL-AA-vWa adopt a binding mode against their host 1TEL domains that buries 269 Å^2^ of solvent-accessible surface area. Specifically, the side chains of Arg 123, Phe 124, Ser 126, Met 129, I 261, and Leu 265 of the vWa domain interacts with the side chains of Tyr 25 and I 25 of the 1TEL domain (Figure 5D). The vWa interface with the vWa domain of an adjacent polymer buries 188 Å^2^ of solvent-accessible surface area. Specifically, the side chain of Gln 138 of the vWa domain interacts with the side chain of Asn 114 of the adjacent vWa domain. Additionally, the side chains of Glu 111 and Asn 107 of the vWa domain interact with the side chains of Pro 144 and Thr 140 of the adjacent vWa domain (Figure 5E). The vWa domain also contacts the 1TEL domain of an adjacent polymer, burying 270 Å^2^ of solvent-accessible surface area. Specifically, the side and main chains of the vWa domain Tyr 208, Val 205, Asp 202, Lue 204 and Tyr 169 contact the side and main chains of the 1TEL domain Leu 11, His 18, and Gln 33, (Figure 5F). The vWa domain also contacts a second vWa domain, burying 144 Å^2^ of solvent accessible surface area. Specifically, the vWa domain Ser 207 side chain makes weak contacts with side chain of Ala 265 in this second vWa domain (Figure 5F).

**Figure 5.**
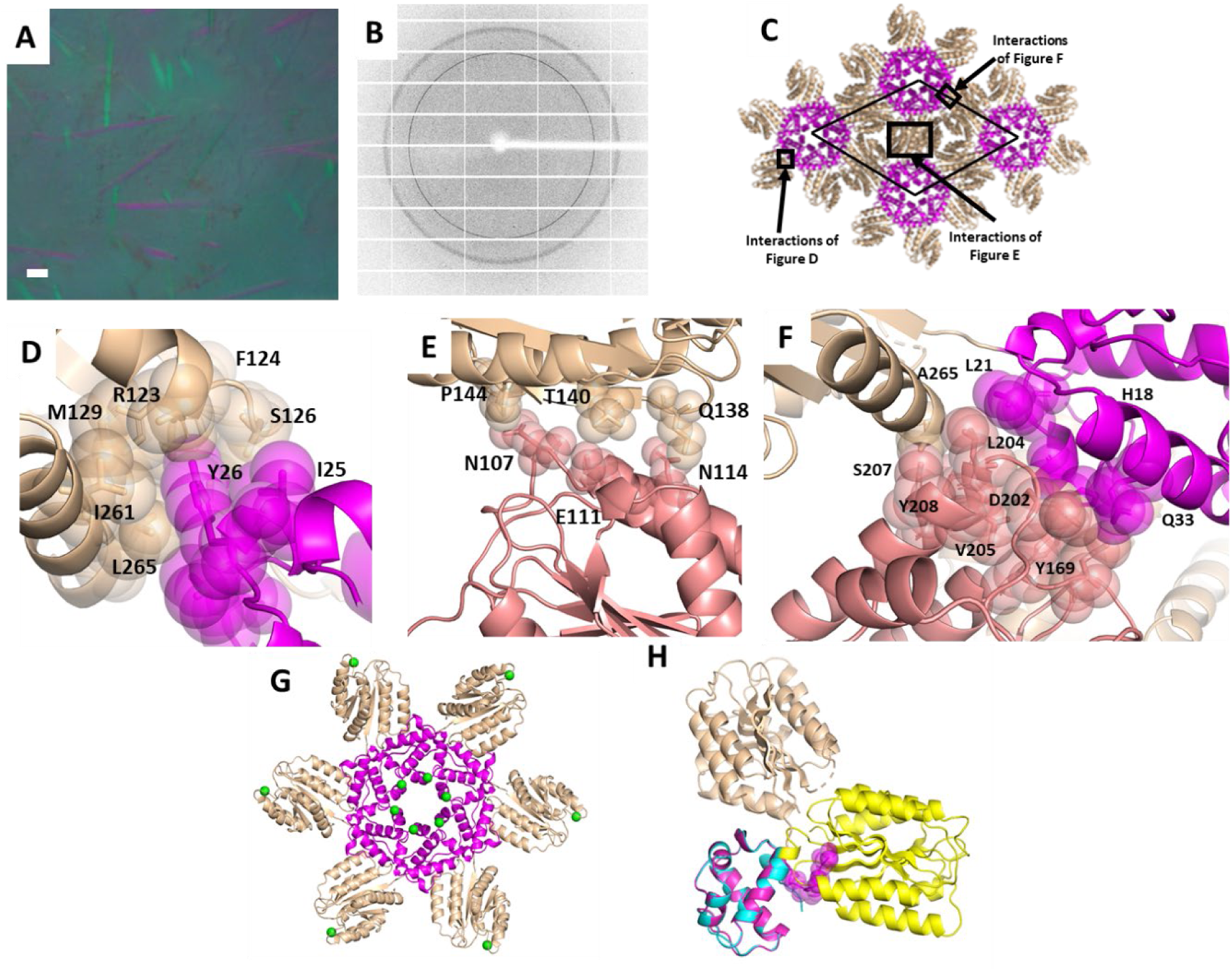
1TEL-AA-vWa crystallization and structure details. A. Representative crystals of 1TEL-AA-vWa. The scale bar is 100 µm. B. Representative diffraction pattern from a 1TEL-AA-vWa crystal. C. Unit cell packing of a 1TEL-AA-vWa crystal with the 1TEL domain in magenta and the vWa domain in wheat. The unit cell is indicated with a thin black line. D. Detail of the interaction between a vWa domain (wheat) and its host 1TEL domain (magenta). The protein is shown in cartoon representation with selected side chains shown as sticks and transparent spheres. E. As in D, but of the interaction between the vWa domain (salmon) and a vWa domain (wheat) from an adjacent TELSAM polymer. F. As in D and E, but highlighting the contacts made by a vWa domain to a second vWa domain and its host 1TEL domain, G. As in D, but highlighting the position of the iodinated tyrosine residues (small green spheres) relative to the 1TEL–vWa polymer. H. As in D, but highlighting the N-terminal 10xHis tag of the 1TEL domain (two histidine residues were resolved in the electron density). The structure of chain A of the 1TEL-TV-vWA (SUMO) structure is superimposed via its 1TEL domain and shown in cyan (1TEL) and yellow (vWa).

Prior to freezing, this crystal was briefly exposed to iodine vapor (Miyatake *et al*., 2006) to label exposed tyrosine side chains. Our object was to assess the degree and location of potential iodine labelling sites on 1TEL for use with target proteins for which molecular replacement is unsuccessful. We observed 79% occupancy iodine labelling of only the less solvent exposed Cε carbon of Tyr 69 of the 1TEL domain, which faces into the solvent void at the center of the 1TEL polymer (Figure 5G). We also observed 64% iodine labeling of only one of the Cε carbons of Tyr 208 of the vWa domain, which makes direct contact to the 1TEL domain of an adjacent 1TEL polymer (Figure 5G). We were also able to resolve the main chain atoms of two of the histidine residues of the 10xHis tag. These histidine residues pack loosely against the main chains of Arg 149, and Ser 153 of the vWa domain from an adjacent 1TEL polymer. Together with the fact that inclusion of the 10xHis tag did not appear to negatively impact crystal nucleation or growth, this suggests that the His tag does not interfere with 1TEL polymerization or target proteins docking to their host polymers. Notably, the vWa domains adopt a binding mode against their 1TEL polymers distinct from that observed with any previous 1TEL--vWa construct, possibly due to the His tag favoring this binding mode or disfavoring the previously-observed binding modes. To test this, we superimposed our previous 1TEL-TV-vWA (SUMO) chain A structure with this 1TEL-AA-vWa structure through their 1TEL domains. This superposition revealed that the His tag occupies the same space as that occupied by the previous 1TEL-TV-vWa (SUMO) structure’s vWa domain, thus forcing the 1TEL-AA-vWa’s vWa domain into an alternative position relative to its host 1TEL domain (Figure 5H). In the light of our prior hypothesis that the 1TEL-TV-vWa (SUMO) construct’s vWa binding mode may be the cause of the polymer-flipping pathology, this provides a possible mechanism for why omission of the cleavable SUMO tag from the 1TEL–vWa constructs prevents these pathologies: The presence of the non-cleavable 10xHis tag blocks access to that vWa binding mode which allows equally energetic parallel and antiparallel polymer-polymer association (and thus polymer flipping).

### 2.4. A more rigid TESAM--target linker did not alter the crystal packing interactions relative to a more flexible linker but did improve the diffraction resolution of 1TEL--vWa crystals

1TEL-TV-vWa formed around 400 crystals (Figure 6A) which is the equivalent to the number of crystals formed by 1TEL-AV-vWa (SUMO) and 1TEL-TV-vWa (SUMO) and higher than 1TEL-AA-vWa (SUMO) (PDB ID: 7N1O) and 1TEL-AA-vWa. 1TEL-TV-vWa formed crystals with an average resolution of 2.0 Å (1.6-2.6 Å across 16 crystals), significantly higher than that of 1TEL-TV-vWa (SUMO), which was 2.7 Å (2.3-3.0 Å), and the highest of all TELSAM fusions tested in this study. Diffracted crystals of 1TEL-TV-vWa indexed an average of 77% of non-ice reflections (31-97%), exhibited an average mosaicity of 0.25° (0.12–0.40°), and had an average ISa of 25 (11–42) (Figure 6B, Table 1). In a subsequent crystallization concentration study, crystals of 1TEL-TV-vWa were observed at concentrations as low as 2 mg/mL and at pH values between 5.0 and 8.5. Notably, this second batch of 1TEL-TV-vWa by the same research team also exhibited no signs of tNCS, twinning, or TELSAM polymer flipping.

**Figure 6.**
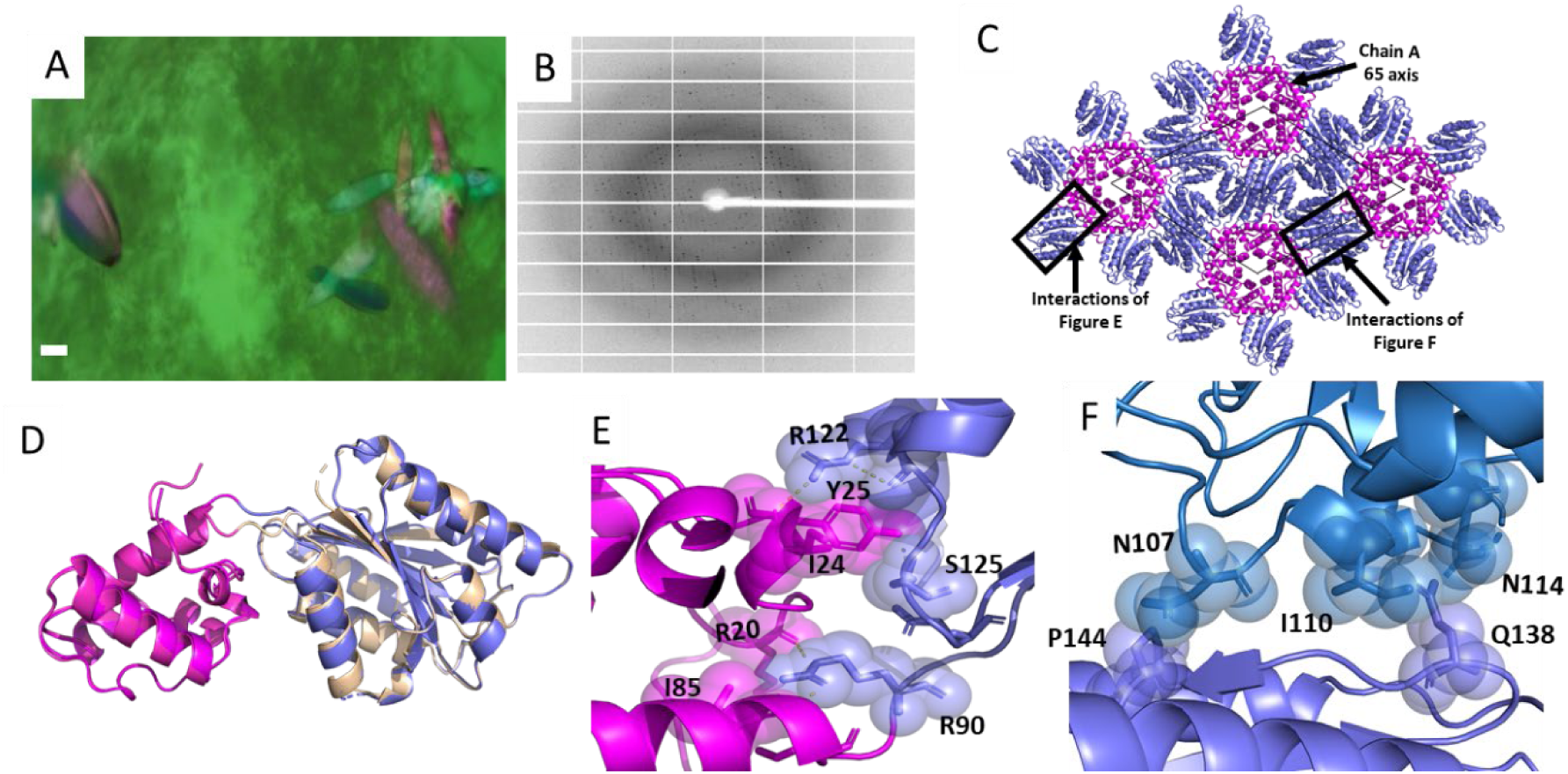
1TEL-TV-vWa crystallization and structure details. A. Representative crystals of 1TEL-TV-vWa. The scale bar is 100 µm. B. Representative diffraction pattern from a 1TEL-TV-vWa crystal. C. Unit cell packing of a 1TEL-TV-vWa crystal with the 1TEL domain in magenta and the vWa domain in purple. The unit cell is shown as a thin black line. D. Superposition of the structure of 1TEL-AA-vWa (magenta and wheat) with the structure of 1TEL-TV-vWa (magenta and slate) through their 1TEL domains. E. Detail of the interaction between the vWa domain (slate) and its host 1TEL domain (magenta). The protein is shown in cartoon representation with selected side chains shown as sticks and transparent spheres. F. As in E, but of the interaction between the vWa domain (slate) and a vWa (blue) domain from an adjacent polymer.

Molecular replacement of 1TEL-TV-vWa was carried out as above, yielding a solution with 1TEL polymers along the 6-fold axes of the P65 unit cell with a = b = 100.9 Å, c = 49.8 Å (Figure 6C). The vWa binding modes against their host 1TEL polymers and the overall crystal packing were very similar to that observed with the 1TEL-AA-vWa construct (Figure 6D). For example, the vWa domains of 1TEL-TV-vWa adopt a binding mode against their host 1TEL domains that buries 470 Å^2^ of solvent-accessible surface area. Specifically, the side chain of Arg 122 and Ser 125 of the vWa domain makes polar interactions with the side chains of Tyr 25 of the 1TEL domain. Arg 90 of the vWa domain also makes polar interactions with Arg 20 and Ile 85 of the 1TEL domain. Unlike 1TEL-AA-vWa, 1TEL-TV-vWa additionally makes many polar interactions between the vWa domain and its own 1TEL domain which presumably make the vWa domain more stable, leading to improved diffraction resolution (Figure 6E). The vWa domain makes glancing contacts to a second vWa domain burying 240 Å^2^ of solvent-accessible surface area. Specifically, the side chain of Gln 138 of the vWa domain forms van der Waals contact with the side chains of Ile 110 and Asn 114 of the adjacent vWa domain. Pro 144 of the vWa domain also interacts weakly with Asn 107 of the adjacent vWa domain (Figure 6F).

We compared the helical rise from the 1TEL-AA-vWa (SUMO), 1TEL-AV-vWa (SUMO), 1TEL-TV-vWa (SUMO), 1TEL-AA-vWa, and 1TEL-TV-vWa crystals to identify relationships between helical rise, linker composition, SUMO inclusion, and diffraction resolution. We observed that the helical rise decreased with increasing average bulk of the amino acids present in the 1TEL–vWa connection (postulated to correlate with rigidity of the linker) in the case of 1TEL-AA-vWa (SUMO), 1TEL-AV-vWa (SUMO), and 1TEL-TV-vWa (SUMO). The same trend was also seen in the case of 1TEL-AA-vWa and 1TEL-TV-vWa. We observed that constructs utilizing a SUMO tag had higher helical rises than constructs without SUMO tag fusion (Figure 7A). The decrease in helical rise also correlates with an improvement in diffraction resolution, which is to be expected. Increased unit cell compactness and lower solvent content would be expected to correlate with increased intermolecular contacts and better resolution (Figure 7A, B) (Table 1).

**Figure 7.**
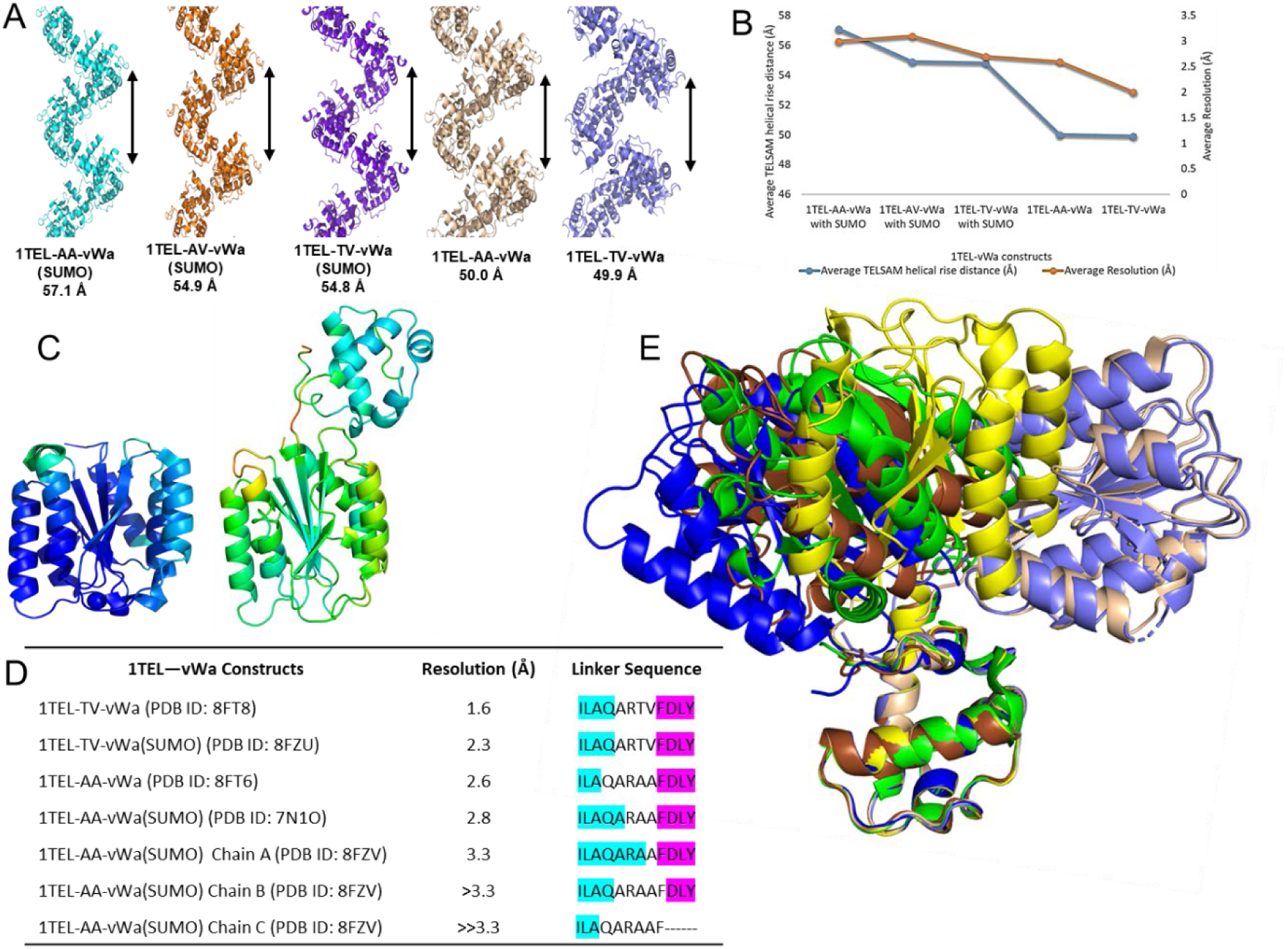
Comparison of structures analyzed in this study. A. Schematic of average helical rise of the 1TEL–vWa constructs discussed in this study. The relative rise of a single turn of each helix is denoted with a black arrow. Fused vWa domains are not shown. B. Plot of the average resolution (orange line) and helical rise (blue line) of the constructs described in this study. C. PDB ID: 1SHU (1.5 Å resolution, left) and 1TEL-TV-vWa (1.6 Å resolution, right) are shown in cartoon representation and colored according to the crystallographic B-factor. The B-factor color scale is the same between the two images. D. Superposition of Batch 1 1TEL-AA-vWa (SUMO) (green), Batch 2 1TEL-AA-vWa (SUMO) chain A (yellow) and chain B (brown), 1TEL-TV-vWA (SUMO) (blue), 1TEL-AA-vWa (wheat), and 1TEL-TV-vWa (slate) through their 1TEL domains. E. Linker sequences for the constructs used in this study. Residues in the 1TEL C-terminal α-helix are highlighted cyan, residues in the vWa N-terminal β-sheet are highlighted magenta, and residues in neither of these secondary structures are uncolored.

1TEL-TV-vWa has comparable diffraction resolution (1.6 Å) to that of the previous highest-resolution structure of the vWa domain, PDB ID: 1SHU (1.5 Å). This is particularly notable because in our hands, crystals of the vWa alone diffract to an average resolution of 2.4 Å (1.9-3.1 Å across 18 crystals), index an average of 56% (19-95%) of non-ice reflections, exhibit an average mosaicity of 0.30° (0.20-0.45°), and an average ISa of 38 (23 - 53). For comparison, we solved and refined the structure of one of these vWa-alone constructs at 2.19 Å resolution (PDB ID: 8FZ4). Data processing and refinement statistics for this structure are given in Table 2. This observation suggests that fusion to 1TEL improved the resolution of vWa crystals produced in our hands by an average of 0.4 Å, a significant improvement (Table 1), providing an exciting hint into what more experienced crystallographers might be able to accomplish with TELSAM mediated crystallization, such as in a recent stunning example involving the SARS Cov2 nsp14 N7-MethylTransferase domain (Kottur *et al*., 2022).

Having a 1.6 Å resolution 1TEL-vWa fusion structure allowed us to directly compare the refined B-factors between this structure and the previous best-resolution structure. PDB ID: 1SHU (Lacy *et al*., 2004) had an average B-factor of 21.1 Å^2^, (0.0-65.9 Å^2^) across just the protein atoms. The vWa domain in the 1TEL-TV-vWa construct had an average B-factor of 46.6 Å^2^ (22.8-97.2 Å^2^), while the 1TEL domain had an average of 35.1 Å^2^ (18.4-96.3 Å^2^) and the linker between them had an average of 60.4 Å^2^ (35.3-99.6 Å^2^) (Figure 7C). This result confirms our earlier hypothesis (Nawarathnage *et al*., 2022) that fusion to 1TEL allows the formation of crystals with comparable diffraction resolution but increased molecular motion in the crystal lattice.

We sought a structural explanation for how increasing the side chain bulk of linker amino acids might improve the resolution of 1TEL–vWa constructs. Increasing the side chain bulk of linker amino acids from Ala-Ala to Thr-Val correlated with a modest improvement of the resolution of 1TEL–vWa (SUMO) and 1TEL–vWa constructs, by 0.3 Å and 0.6 Å, respectively. Our structures reveal that substitutions to the amino acids in the linker do not make it fully rigid, but rather appear to reduce the number of highly flexible residues present. We note that in all 1TEL–vWa structures solved to date, there is always at least one amino acid in the linker that is not part of the 1TEL C-terminal α-helix or the vWa N-terminal β-sheet, such as in Batch 1 (PDB ID: 7N1O) (Nawarathnage *et al*., 2022) or in the Batch 2 1TEL-AA-vWa (SUMO) chain A (Figure 7D). This observation suggests that a fully rigid connection between 1TEL and a target protein may not be favorable for crystal formation, as we suggested previously. This is a property that we are testing in more depth in a follow-up study. Alignment of the 1TEL domains of all 1TEL–vWa structures solved to date reveals that there is some correlation between the number of linker residues not part of the 1TEL C-terminal α-helix or the vWa N-terminal β-sheet and the observed or extrapolated (from the resolution and lack of crystal defects) order of the fused vWa domain (Figure 7E). For example, 1TEL-TV-vWa had four such amino acids and yielded a 1.6 Å resolution structure, while 1TEL-AA-vWa had five such amino acids and yielded a 2.6 Å structure. Likewise in the Batch 2 1TEL-AA-vWa structure, chain A had one linker amino acid not in a regular secondary structure and had a resolution < 3.3 Å, while chain B had five such linker amino acids and a resolution ∼3.3 Å, and chain C had 6 such linker amino acids and a resolution of >> 3.3 Å. We further note that in structures of 1TEL-TV-vWa constructs, with or without SUMO, the valine side chain of the Thr-Val linker segment is nearly always well-resolved in the electron density, while the threonine never is, suggesting that it adopts a number of side chain conformational states. This fits well with the 1TEL-AV-vWa (SUMO) linker variant crystallizing nearly as well as the 1TEL-TV-vWa (SUMO) variant while the 1TEL-TT-vWa (SUMO) variant did not form crystals.

The 1TEL–vWa constructs characterized to date reveal that the vWa domain can adopt at least nine distinct binding modes against their host polymers, four of them clearly observed and five of them extrapolated from partial electron density (Figure 7E). Interestingly, the choice of vWa binding mode showed no correlation to the type of linker used (Figure 7D) but was strongly correlated with the presence or absence of the SUMO tag and/or the batch of protein. Curiously, 1TEL-AV-vWa (SUMO) (to be described elsewhere) and 1TEL-TV-vWa (SUMO) were prepared in separate batches by the same research team, but exhibited the same vWa binding mode. Likewise, 1TEL-AA-vWa and 1TEL-TV-vWa were prepared in separate batches by this same research team but also exhibited the same vWa binding mode, distinct from that of 1TEL-AV-vWa (SUMO) and 1TEL-TV-vWa (SUMO). Taken together, these observations highlight the still very significant factor of batch-to-batch variation and researcher skill level on crystallization dynamics and thus on crystallization propensity and quality, regardless of whether or not TELSAM fusion is employed.

## 3. Discussion

Fusion to TELSAM has now been demonstrated in multiple cases to allow the rapid crystallization of even difficult target proteins (Kottur *et al*., 2022), to enable the achievement of high-resolution data collection (<2.0 Å), to allow crystallization at unusually low protein concentrations, and to result in crystals where the protein of interest has an increased degree of residual molecular motion, as evidenced by significantly higher crystallographic B-factors. These higher B-factors hint that TELSAM-mediated crystals may capture proteins of interest in subtly more physiological conformations, with less interference from crystal packing artefacts. If this is true, it may open the door to higher-accuracy studies of protein structure and dynamics, potentially benefitting fields such as protein and enzyme engineering and the study of intrinsically disordered and partially folded proteins. Due to TELSAM’s built-in validations (TELSAM subunits most often form the expected six-fold helical polymer and the target proteins must be with an appropriate distance of the TELSAM C-terminus) TELSAM-fusion crystals are a promising system to enable the phase solution and structure determination of such difficult proteins.

In replicating our previous results, we identified and investigated several parameters unexplored in our initial pilot study (Nawarathnage *et al*., 2022), as well as previously unencountered issues, such as target proteins that are not fully resolved in the resulting electron density. To better understand the best practices for TELSAM-mediated crystallization and improve the reliably of the technique, we experimented with modifications to the 1TEL–vWa linker and found that doing so modestly improved diffraction resolution and appeared to enforce a single vWa domain binding mode to the host 1TEL polymer. Based on current evidence, we propose that the optimal linker between 1TEL and a protein of interest consists of a small number (∼2) of flexible residues and places the protein of interest as close to the 1TEL polymer as possible.

The fact that three out of the four linkers tested in this study resulted in crystallization and that some linkers clearly outperformed others suggests the utility of designing and testing a number of different linker variants to optimize the chances of crystallization and of obtaining a high-resolution (< 2.0 Å) structure of a protein of interest. This parallels the experience of Kottur et al., where three different linkers were tested in an attempt to crystallize the recalcitrant SARS Cov2 nsp14 N7-MethylTransferase. In this case, only one of the three linkers resulted in crystals, but with optimization of the crystallization conditions, the authors were able to reliably achieve 1.4-1.6 Å diffraction resolution (Kottur *et al*., 2022).

We identified the phenomenon of TELSAM polymer flipping and were able to largely correct for it by differently scaling the weak and strong reflections in our data. The approach used to correct the data could provide a way forward for other systems that exhibit similar diffraction data pathologies. We observed that polymer flipping appears to correlate with the specific binding mode of the target protein to its host TELSAM polymer. Based on this observation, we hypothesize that polymer flipping is possible whenever the fused protein of interest adopts a binding mode to its host TELSAM polymer that allows parallel and antiparallel TELSAM polymer interactions with roughly similar binding energy. While inclusion of a non-cleavable 10xHis tag appears to have blocked the vWa binding mode associated with TELSAM polymer flipping in the current case, we expect that polymer flipping may be dependent on the specific size, shape, and preferred binding mode of the target protein. This phenomenon and its effective treatment are the subject of ongoing study.

Based on the extremely minimal contacts observed between the vWa domain and its host or adjacent 1TEL polymers, we hypothesize that target proteins choose the binding mode to their host TELSAM polymers at the time of polymer-polymer association. We also hypothesize that to obtain a high-resolution structure with a well-resolved target protein, all target proteins must adopt the same binding modes to their host polymers throughout the crystal and that TELSAM–target polymers must then adopt a single binding mode one to another. The somewhat lower fraction of reflections indexed in some TELSAM-fusion crystals may hint at failure to completely achieve these two requirements throughout the entirety of these crystals, although this has not prevented high-resolution structure solution in the cases reported here.

We provide evidence that trace protease activity can cleave the target protein from the 1TEL chaperone and that this can result in unusual TELSAM crystallization behaviour, such as a lack of canonical TELSAM polymer formation. Other instances of target protein cleavage have resulted in crystals of only 1TEL polymers. We have also observed other phenomena that appear to correlate with cleavage of the target protein by trace proteases, including TELSAM double helices and TELSAM polymer compression. These will be described elsewhere. At present, the connection between SUMO tag inclusion and these outcomes is unconfirmed, as are the factors that direct a given crystallization experiment toward one outcome or another.

Based on these observations, we currently recommend omitting cleavable fusion tags from TELSAM-target protein fusion constructs because of the demonstrated behavior of SUMO protease or another co-purified protease in cleaving target proteins from TELSAM polymers. Additionally, retaining the 10xHis tag during crystallization clearly does not block crystal formation and may improve order and resolution by limiting the possible binding modes of the target protein to its host TELSAM polymer.

## 4. Materials and methods

### 4.1. Cloning of 1TEL-AA-vWa (SUMO), 1TEL-AV-vWa (SUMO), 1TEL-TV-vWa (SUMO), 1TEL-TT-vWa (SUMO), 1TEL-AA-vWa, and 1TEL-TV-vWa

The 1TEL-AA-vWa (formerly 1TEL-flex-vWa) construct from Nawarathange et al. was used as the starting point for the design of the 1TEL—vWa variants used in this study. This construct placed residues 40–217 of the human Anthrax toxin receptor 2 vWa domain (ANTXR2, also known as capillary morphogenesis gene 2, CMG2, Uniprot: P58335) downstream of residues 47–123 (the SAM domain) of human Transcription Factor ETV6 (ETS variant transcription factor 6, also known as translocation ETS leukemia, TEL, Uniprot: P41212). The pH-sensitive variant of this SAM domain is hereafter referred to as 1TEL. An alanine linker was placed between the two domains and substitutions to relative to the wild type proteins included R49A, V112E, K122A, R41A, and C175A (Nawarathnage *et al.,* 2022). The AV, TV, and TT substitutions were designed at positions 41 and 42 of the vWa domain in PyMOL (The PyMOL Molecular Graphics System, Version 2.0 Schrödinger, LLC) and Geneious (Geneious 9.1.8, https://www.geneious.com). Gene fragments were synthesized by Twist Bioscience (www.twistbioscience.com) and were assembled into the pET42_SUMO vector (Walls *et al*., 2022) (cut with XhoI) using Gibson cloning (Gibson *et al*., 2009) to generate 1TEL-AV-vWa (SUMO), 1TEL-TV-vWa (SUMO), and 1TEL-TT-vWa (SUMO) constructs. The 10xHis-1TEL-AA-vWa and 10xHis-1TEL-TV-vWa were cloned in the same manner, being assembled into a pET42_SUMO vector first cut with XhoI and NdeI to remove the SUMO domain. The sequence MGHHHHHHHHHH was appended to the N-terminus of 1TEL for these SUMO tag-free constructs. All constructs were introduced into BL21(DE3) cells and sequence verified by Sanger sequencing in both directions by Eton Bioscience (www.etonbio.com).

### 4.2. Cloning of 1TEL-G-Ack1-UBA

Residues 958-1038 (the UBA domain) of human Activated CDC42 kinase 1 (Ack1, also known as Thirty-eight Negative Kinase-2 or Non-receptor tyrosine-protein kinase TNK2, Uniprot: Q07912) were reverse-translated, codon-optimized using DNAworks (Hoover & Lubkowski, 2002) and placed downstream of 1TEL, separated by a single glycine linker. The sequence MGHHHHHHHHHH was appended to the N-terminus of 1TEL. This construct included an R80S substitution (carried over from the structure of the TELSAM domain used in the modeling (PDB ID: 2QAR,(Nauli *et al*., 2007)). This construct was synthesized by a commercial vendor, assembled into the pET42-SUMO, previously cut with XhoI and NdeI (Walls *et al*., 2022), introduced into BL21(DE3) cells, and sequence verified as above.

### 4.3. Cloning of 1TEL-helix-TNK1-UBA

Residues 590-666 of human Thirty-eight Negative Kinase-1, also known as Non-receptor tyrosine-protein kinase TNK1, Uniprot: Q13470) were reverse-translated, codon-optimized, and placed downstream of 1TEL, separated by a linker with the sequence RDLE, intended to form a continuous α-helix between the 1TEL C-terminus and the UBA N-terminus. Both cysteines in the UBA domain (C610 and C644) were mutated to alanines to prevent oxidation-mediated misfolding. The construct was synthesized by a commercial vendor, assembled into the pET42-SUMO, previously cut with just XhoI (Walls *et al*., 2022), introduced into BL21(DE3) cells, and sequence verified as above.

### 4.4. Protein expression

20 ml of Luria–Bertani (LB) medium with 100 µg ml^−1^ kanamycin was inoculated with a stab of frozen cell stock and shaken at 37 °C and 250 rpm overnight. The following day, 15 ml of the overnight culture was added into 1 L of LB medium supplemented with 0.05% glucose and 100 µg ml^−1^ kanamycin and incubated at 37 °C and 250 rpm until the optical density (OD) reached 0.6. At OD 0.6, isopropyl β-D-1-thiogalactopyranoside (IPTG) was added to a final concentration of 100 µM. The culture was incubated at 18°C and 200 rpm overnight. The following day, the cells were collected by centrifugation, snap-frozen in liquid nitrogen, and stored at −80°C.

### 4.5. Purification of1TEL-AA-vWa (SUMO),1TEL-AV-vWa (SUMO),1TEL-TV-vWa (SUMO), and 1TEL-TT-vWa (SUMO)

All purification steps were performed on ice or in a 4°C refrigerator. Cell pallets (20 g) were resuspended in a five-fold excess of wash buffer (50 mM Tris, 200 mM KCl, 50 mM imidazole pH 8.8) containing 1 mM phenylmethylsulfonyl fluoride (PMSF), 100 µM dithiothreitol (DTT), 0.5 mg/mL lysozyme, and 800 nM deoxyribonuclease I. Suspended cells were then sonicated at 60% power with 12s on/59 s off for 25 cycles (Qsonica Q500) in a spinning ice bath. The resulting lysate was centrifuged at 40,000 g and supernatant was loaded onto 6 ml of HisPure Ni-NTA resin (Thermo Scientific). The column was then washed with 7-column bed volumes (CV) of wash buffer (50 mM Tris, pH 8.8, 200 mM KCl, 50 mM imidazole). The protein was then eluted with about 7 CV of elution buffer (50 mM Tris, pH 8.8, 200 mM KCl, 400 mM imidazole) until protein stopped appearing in subsequence fractions, as detected with Bradford reagent (Bradford, 1976). The collected protein was then desalted using several PD-10 desalting columns in parallel (Cytiva). The SUMO tag was removed by incubating the protein overnight at 4°C with 0.5 mg SUMO protease per 100mL of protein. The SUMO protease and cleaved SUMO tags were removed by flowing the cleavage reaction over 2 ml of fresh Ni-NTA resin. The protein was then concentrated to 3 mL and diluted ten-fold with water (to decrease the conductivity below 3 mS/cm). The diluted protein was then loaded onto 5 ml Source 15Q anion exchange resin (Cytiva) and eluted with 50 mM Tris, pH 8.8, 1M KCL using a gradient elution. Fractions containing the pure protein of interest were concentrated to 2 mL and loaded onto a 100 ml Superdex 200 Prep Grade size exclusion column (Cytiva). The proteins were judged to be greater than 95% pure by SDS-PAGE.

### 4.6. Purification of 1TEL-AA-vWa, 1TEL-TV-vWa, 1TEL-G-Ack1 and 1TEL-helix-TNK1-UBA and vWa alone

These proteins were purified in an identical manner to their SUMO-tagged counterparts above, except that the SUMO tag cleave, tag removal, and ion exchange chromatography steps was omitted.

### 4.7. Crystallization and diffraction of 1TEL-AA-vWa (SUMO),1TEL-AV-vWa (SUMO),1TEL-TV-vWa (SUMO),1TEL-TT-vWa (SUMO), 1TEL-AA-vWa, and 1TEL-TV-vWa and vWa alone

The concentration of the purified protein was adjusted to 1 and 20 mg/mL. 1TEL-TV-vWa was additionally adjusted to 0.1, 0.2, 0.5, 1, 2, 5, 10, 15, and 20 mg/mL for the protein concentration trial. The vWa alone concentration was adjusted to 20 mg/mL. Commercially available crystallization screens (PEG Ion, Salt RX, and Index (Hampton Research)) and custom screens (PEG-custom and Ammonium Sulfate) were screened at the above-given protein concentrations. 1 μL of the indicated protein solutions were combined with 1 μL of reservoir solution (SPT Labtech Mosquito) and equilibrated against 50 μL of reservoir solution in a sitting drop vapor diffusion format. Crystals appeared after 2 days for 1TEL-TV-vWa, 3-7 days for1TEL-AA-vWa (SUMO), 1TEL-AV-vWa (SUMO), 1TEL-TV-vWa (SUMO), and 1TEL-AA-vWa, 35 days for the vWa alone. No crystals appeared for 1TEL-TT-vWa (SUMO). The largest 1TEL-AA-vWa (SUMO) crystals appeared in 100 mM Bis-Tris, pH 5.7, 3.0 M NaCl. The largest 1TEL-AA-vWa crystals appeared in 100 mM Bis-Tris, pH 5.7, 3.0 M NaCl, and 0.07 M BisTris propane, pH 7.8, 0.03 M citric acid, 20% PEG3350. The largest 1TEL-AV-vWa (SUMO), 1TEL-TV-vWa (SUMO), and 1TEL-TV-vWa crystals appeared in 1.3-2.0 M ammonium sulfate, 0.1M HEPES, pH 5-8.5. Crystals were cryoprotected by briefly passing them through a solution of 20% glycerol in reservoir solution, prior to freezing in liquid nitrogen.

### 4.8. Crystallization 1TEL-G-Ack1 and 1TEL-helix-TNK1-UBA

1.2 μL of 5 mg/mL protein was combined with 1.2 μL of reservoir solution over 50 uL of reservoir solution in a sitting drop format (SPT Labtech Mosquito). Commercially available crystallization screens (PEG Ion and Index, Hampton Research) and custom screens (BisTris Mg-Formate) were used. Crystals appeared in three days in five distinct crystallization conditions. The largest (200 μm) appeared in 100 mM Bis-Tris pH 7.0, 200 mM Mg-formate. Before freezing in liquid nitrogen, the crystals were cryoprotected using 20% glycerol in crystallization reservoir solution.

### 4.9. Data collection, reduction, and structure solution

X-ray diffraction data were collected remotely at SSRL beamlines 9-2, 12-1, and 12-2. Most crystals were diffracted at 12658 eV energy. 1TEL-AV-vWa (SUMO) and selected 1TEL-TV-vWa (SUMO) crystals were diffracted at an energy of 12398 eV. Iodine-treated crystals were diffracted at an energy of 6690 eV. Typically, 120-360° of data were collected in 0.1 or 0.2° increments with 0.2 s exposures.

The Autoproc pipeline (Vonrhein *et al*., 2011) was used to process the datasets, and molecular replacement using Phenix Phaser (Liebschner *et al*., 2019, McCoy *et al*., 2007) was employed to solve the phases. Structure rebuilding was done in Coot (Emsley *et al*., 2010) and refinement in PhenixRefine (Afonine *et al*., 2012, Williams *et al*., 2018). TLS parameters were refined using TLS groups determined by Phenix. Refinement was assisted using statistics from the MolProbity server (Chen *et al*., 2010).

The diffraction images of the 1TEL-TV-vWa (SUMO) crystal were processed with DIALS (Winter *et al*., 2018) using the global background model (Parkhurst *et al*., 2016).The integrated data were scaled and merged using Aimless (Evans & Murshudov, 2013). The complete dataset of intensities was split into two datasets with h-k=3n and h-k=3n+/-1 using a purpose-made script written in R (Team, 2021). The I to F conversion was completed separately for the two partial datasets using ctruncate from the CCP4 suite (Winn *et al*., 2011). The initial model was obtained using the h-k=3n dataset and Molrep (Vagin & Teplyakov, 1997). The h-k=3n and h-k=3n+/-1 datasets were separately rescaled to Fcalc using Refmac (Murshudov *et al*., 2011) with zero cycles of coordinate refinement and joined using an R-script. The atomic model was then refined against the joined dataset using Refmac. The procedure involving separate scaling and joining. Model refinement was repeated four times, alternated with sessions of model inspection and corrections with Coot (Emsley *et al*., 2010). To evaluate the structure amplitude correction that has, in effect, been applied to the h-k=3n+/-1 dataset, the final h-k=3n and h-k=3n+/-1 datasets were fitted separately to the complete set of structure amplitudes generated from the complete set of merged intensities using ctruncate. The fitting was completed using an R script, and the relative values of the overall scale factor, 2.658(2), B11, 5.18(3) Å^2^, and B33, 2.13(4) Å^2^, were calculated from the respective absolute values. This translates into a relative scaling coefficient of about 7 in terms of intensities.

## Acknowledgements

Research reported in this study was supported by the National Institute of General Medical Sciences of the National Institutes of Health under award number R15GM146209. The content is solely the responsibility of the authors and does not necessarily represent the official views of the National Institutes of Health.

Use of the Stanford Synchrotron Radiation Lightsource, SLAC National Accelerator Laboratory, is supported by the U.S. Department of Energy, Office of Science, Office of Basic Energy Sciences under Contract No. DE-AC02-76SF00515. The SSRL Structural Molecular Biology Program is supported by the DOE Office of Biological and Environmental Research, and by the National Institutes of Health, National Institute of General Medical Sciences (P30GM133894). The contents of this publication are solely the responsibility of the authors and do not necessarily represent the official views of NIGMS or NIH.

